# A mammary-specific microfluidic device for studying post-radiotherapy vascular-immune cell interactions

**DOI:** 10.64898/2026.09.03.749257

**Authors:** Shannon E. Martello, Sofia N. Luna, Katelyn Derr, Gladys Martinez Franco, Marissa Paul, Mark Mc Veigh, Leon M. Bellan, Marjan Rafat

## Abstract

Radiotherapy (RT) significantly improves outcomes and reduces the risk of recurrence in breast cancer. However, in patients that experience recurrence despite treatment, the irradiated breast tissue may create a pre-metastatic niche that promotes tumor cell infiltration. Because current models of metastasis lack key physiological and microenvironmental cues, we developed a mammary vasculature-on-chip (MVoC) to model how the vasculature responds to RT and impacts immune and tumor cell behavior in mammary tissue. Murine and human MVoCs incorporated endothelial cells (ECs) cultured under physiologic shear stress and mammary-specific fibroblasts to mimic the vascular-fibrous stroma interface. MVoCs exhibited a transient, acute response to RT that led to a persistent phenotypic shift. Neutrophils, which play key roles in pre-metastatic niche formation, adhered more to irradiated ECs and induced persistent damage to the endothelial barrier, preventing recovery. Neutrophils also significantly increased tumor cell adhesion to the endothelium in MVoCs. These findings demonstrate that MVoCs can be used to study RT-induced pre-metastatic niche formation and suggest a positive feedback loop between damaged ECs and immune cell activation that may facilitate tumor cell colonization. MVoCs provide a biologically relevant approach to probe molecular and cellular interactions, improving translation toward developing novel therapies that improve patient outcomes.

## Introduction

Radiotherapy (RT) remains part of the standard of care for breast cancer patients as it significantly reduces the risk of recurrence^[1]^. Despite this, up to 20% of patients with triple-negative breast cancer, an aggressive subtype that lacks targeted therapies, will still experience treatment failure and recurrence within five years of an initial diagnosis^[2–4]^. RT does not spare the adjacent normal breast tissue and increasing evidence suggests that when failure occurs, this damaged normal tissue exhibits chronic dysfunction that actively contributes to recurrence within the field of RT. Recent pre-clinical studies demonstrate that local remodeling of normal tissue microenvironments post-RT creates a pre-metastatic niche that promotes immune cell recruitment, vascular dysfunction, and extracellular matrix reorganization that ultimately leads to colonization of circulating tumor cells (CTCs) that shed from the primary tumor^[4–8]^. While these studies support the hypothesis of pre-metastatic niche formation in irradiated mammary tissue, the effects of RT on the mammary microenvironment and the mechanisms governing tumor cell seeding at this site remain poorly understood.

The microvasculature serves as a critical barrier between tumor and immune cells in circulation and pre-metastatic niches, making it a key regulator of metastasis^[9,10]^. When circulating cells are attracted to a site, they must interact with the endothelial cells (ECs) lining microvessels, adhering and rolling along the endothelium before extravasating into the tissue. RT is known to cause acute vascular damage by causing EC apoptosis, increasing vessel permeability, and inducing inflammatory signaling that attracts neutrophils^[11]^. Depending on factors like RT dose, fractionation schedule, and a patient’s systemic immune status, this acute response may lead to chronic dysfunction from persistent inflammation and EC senescence that may facilitate a microenvironment fit for CTC infiltration and tumor recurrence^[7,12–14]^. Despite the vasculature being a key modulator of tumor cell seeding and the normal tissue response to RT, most mammary tissue studies focus on the epithelial or fibroblast response to treatment^[15–17]^, leaving the vascular contributions to recurrence post-RT understudied.

Current experimental models provide limited ability for investigation of RT-induced tumor and immune cell trafficking and their interactions with the mammary tissue microenvironment. While *in vivo* models are historically the gold standard for pre-clinical testing and treatment discovery, these models have poor clinical translation, high costs, and lack the ability to dynamically probe and observe cell-cell interactions^[18,19]^. On the contrary, conventional 2D and 3D models (e.g., flasks, transwell models, organoids) enable higher throughput probing of molecular signaling but lack the multicellular organization and physical cues present *in vivo* that are necessary for understanding response to RT^[20–22]^. Microphysiological systems (MPS) and organ-on-a-chip (OoC) models bridge this gap by integrating multiple organ-specific cell types with physiologically relevant mechanical cues to model whole organs, organ-specific functional units, or tissue barrier interfaces, yielding a model that faithfully recapitulates *in situ* cellular responses while enabling isolated analysis of specific cell types^[23–26]^. Although MPS and OoCs have been increasingly used to model vascular dysfunction, inflammation, and tumor cell extravasation[27–31], their application to radiation biology remains relatively limited. Existing radiobiology studies using OoCs have largely focused on tumor models to understand radiosensitivity and identify potential candidates for combination therapy^[32–34]^. Additional studies focus on the normal tissue response to radiation^[35–38]^. However, those studies primarily emphasized organ-specific radiotoxicity and identification of radioprotectors or countermeasures for application to off-target RT effects, accidental exposure in a nuclear disaster, or occupational exposure. Few models have been designed to understand how RT alters the normal tissue vasculature and impacts subsequent pre-metastatic niche formation. In particular, there remains a need for a mammary-specific platform that incorporates physiological shear stress and surrounding stroma to study how RT alters the vascular niche and its interactions with circulating tumor and immune cells.

To address this gap, we developed a mammary vasculature-on-chip (MVoC) model—a perfused OoC model of the mammary vascular-stromal interface—to understand the impact of RT and immune cell infiltration on tumor cell adhesion to the endothelium. Using 3D-printing and photolithography, we fabricated MVoC devices that were used to culture murine and human ECs with mammary-specific fibroblasts embedded in extracellular matrix (ECM) to facilitate the microenvironmental cell signaling at the mammary vascular-stromal interface. We found that when ECs were exposed to RT, MVoCs initially exhibited a typical acute response to radiation with an increase in permeability, followed by recovery of the endothelial barrier and signs of senescence induction as early as 7 days post-RT. Importantly, we found that RT increased neutrophil adhesion in both murine and human models and that pre-conditioning of MVoCs with neutrophils caused increased tumor cell adhesion, demonstrating the potential of our model to evaluate pre-metastatic niche formation and the metastatic cascade in RT-treated mammary tissue.

## Materials and Methods

### Microfluidic Device Fabrication

To model the 3D interface of the vascular-stromal barrier within the mammary gland, we employed a 3-layer design^[39]^, including two vertically-stacked, parallel channels separated by a porous membrane to create a vascular compartment (top channel) and a fibrous mammary stroma compartment (bottom channel). Designs for top and bottom channel molds were created with Fusion 360 (Autodesk). Top channel molds were fabricated using Liqcreate Bio-Med clear resin (Formlabs) and an mSLA 3D Sonic Mini 8Ks printer (Phrozen) with 22 µm, 22 µm, and 50 µm resolutions across the *x*-, *y*-, and *z*-axes, respectively. Prints were washed by sonicating in 70% ethanol until visibly clean and compressed air was used to remove excess resin. The print was then cured on an Anycubic Wash&Cure 2.0 (Anycubic) for 30 min. Cured top channel prints were coated with 2 g of parylene-C using a Labcoter PDS 2010 parylene deposition machine (Specialty Coating Systems)^[40]^. Bottom channel molds were fabricated using SU8 XFT100 photoresist (Kayaku Advanced Materials) and standard photolithography techniques. Final feature heights of both top and bottom channel molds were measured using a stylus profilometer (KLA Tencor P-7).

Top and bottom channel layers were then fabricated using soft lithography. Briefly, poly(dimethylsiloxane) (PDMS; Sylgard 184, Dow) base polymer was mixed at a 1:10 w/w ratio with the curing agent and poured into the top (25 g) and bottom (5 g) channel molds. After degassing under vacuum to remove bubbles, layers were cured for 4 hr at 65°C. PDMS slabs were demolded and microchannel features were cut into approximately 3 mm x 4 mm slabs. Inlet ports were punched in top channel layers using 1 mm biopsy punches (Miltex). Layers were taped to cover microchannels and ports and stored at room temperature until device assembly.

Poly(ethylene terephthalate) (PET) membranes (3 µm pores; Sterlitech) were cut into 1 mm x 3 mm rectangles and sonicated in isopropyl alcohol (IPA) for 5 min to clean. Membranes were then surface-modified with poly[dimethylsiloxane-co-(3-aminopropyl)methylsiloxane] (amine-PDMS linker; Sigma) by coating each side with the linker and baking at 80°C for 1 hr per side. Excess linker was removed by sonicating membranes in IPA for 1 min.

To assemble devices, layers were treated with oxygen plasma (Harrick Plasma PDC-32G) for 30 sec and brought into contact to facilitate irreversible bonding. This was done in a three-step process to bond: (1) top channel layer and membrane, (2) top channel/membrane construct and bottom channel layer, and (3) assembled device and glass slide. Final devices were incubated for at least 12 hr at 65°C to ensure complete bonding between all layers. Fine-point tweezers were used to puncture a hole in the membrane of bottom channel inlet and outlet ports to allow fluid flow through the bottom channel. Devices were sterilized by flowing 70% ethanol through each channel and exposing to UV light for 30 min. Sterilized devices were dried and stored at RT until use.

### Mammary Stromal Vascular Fraction Isolation

Primary cells were harvested from mice in accordance with institutional guidelines and protocols approved by the Vanderbilt University Institutional Animal Care and Use Committee. Isolation of cells from the mammary stromal vascular fraction (SVF) was performed as previously described^[16]^. Briefly, female BALB/c mice aged 6-8 weeks old were euthanized in a CO_2_ chamber with cervical dislocation as a secondary physical method as required by the NIH Office of Laboratory Animal Welfare. Inguinal MFPs were immediately harvested, minced, and digested in 212.5 µg/mL liberase (Roche) for 45 min at 37°C with agitation. MFPs were then filtered using 100 µm filters before plating into two 6 cm dishes per mouse. Cells were expanded at 5-7 days (passage 1) and 8-10 days (passage 2) post-isolation before using in MVoCs. Media was changed every other day.

### Cell Culture

For murine MVoCs, murine embryonic ECs (C166; ATCC, CRL-2581) were cultured in Dulbecco’s Modified Eagle Medium (DMEM; Gibco) supplemented with 1% penicillin/streptomycin and 10% fetal bovine serum (FBS). Primary murine SVF was cultured in DMEM supplemented with 1% penicillin/streptomycin and 20% bovine calf serum (BCS). Murine neutrophils were differentiated from the murine promyelocyte cell line (MPRO 2.1; ATCC CRL-3604) by supplementing base media with 10 µM all-trans retinoic acid (ATRA; Sigma) for at least 72 hr. Base media for MPRO consisted of Iscove’s Modified Dulbecco’s Medium (IMDM; Gibco) supplemented with 1% penicillin/streptomycin, 20% horse serum, and 10 ng/mL granulocyte-macrophage colony-stimulating factor (GM-CSF). dTomato-labeled 4T1 cells (4T1-dTom) were obtained from Dr. Laura Bronsart (Stanford University) in 2017. For human MVoCs, human umbilical vein endothelial cells (HUVECs; ATCC, PCS-100-013) were cultured in EGM-2 with supplement mix provided by the manufacturer (PromoCell). Human mammary fibroblasts were originally isolated from reduction mammoplasty tissue^[41]^ and were maintained in DMEM supplemented with 1% penicillin/streptomycin and 10% BCS. For human neutrophil experiments, HL-60 cells (ATCC, CCL-260) were differentiated by supplementing base media (DMEM/20% FBS/1% penicillin/streptomycin) with 1.25% dimethyl sulfoxide and 1 µM retinoic acid and culturing for 5 days. MDA-MB-231 cells (ATCC, HTB-26) were maintained in DMEM supplemented with 1% penicillin/streptomycin and 10% FBS. All cells were cultured in a humified incubator at 37°C with 5% CO_2_ and tested monthly for mycoplasma using a MycoAlert Mycoplasma Detection Kit (Lonza).

### Flow Cytometry

SVF and RMFs were collected via trypsinizing to create a single cell suspension. Cells were stained with Aqua fixable viability stain (Invitrogen) and FC receptors were blocked with CD16/32 (Biolegend) while simultaneously staining for all surface markers. Cells were incubated at 4°C for 20 min and fixed at least overnight in 1% neutral buffered formalin. For intracellular staining, cells were permeabilized for 5 min in permeabilization buffer (eBioscience), washed, and stained for intracellular proteins for 30 min at room temperature. Flow cytometry was then performed on a three-laser Amnis CellStream machine (Cytek). FlowJo (Becton Dickinson) was used for analysis. The following antibodies were used for surface marker analysis: CD45-BV650 [30-F11] (1:200; BioLegend 103151, lot no. B410647); CD31-FITC [390] (1:100; eBioscience 11-0311-81, lot no. 2191638); PDGFR -SuperBright 600 [APA5] (1:300; eBioscience 63-1401-82, lot no. 2255498); PDGFRβ-APC-eFluor780 [APB5] (1:100; eBioscience 47-1402-82, lot no. 2427257); Thy1.2/CD90.2-APC [53-2.1] (1:100; eBioscience 17-0902-82, lot no. 2213093); Podoplanin-PE-Cy7 [8.1.1] (1:100; eBioscience, 25-5381-82, lot no. 2416193); CD26-BV421 [H194-112] (1:100; BD 740021, lot no. 1343660); and fibroblast activation protein-AF750 (FAP) [983802] (1:200; Novus FAB3715S, lot no. 1664396). The following antibodies were used for intracellular analysis: pan-cytokeratin-APC (PanCK) [C-11] (1:100; Invitrogen MA1-10325, lot no. WH3366013); E-cadherin-PerCP-eFluor710 [DECMA-1] (1:200; eBioscience 46-3249-82, lot no. 2148136); and Vimentin-APC [280618] (1:100; Novus IC2105A, lot no. XB3513219).

### Mammary Vasculature-on-chip (MVoC) Culture

To facilitate ECM binding to device membranes and improve cell attachment, sterilized devices were treated with 0.5 mg/mL sulfo-SANPAH (Thermo Scientific), activated by a UV box over two 10 min incubations. Channels were then rinsed with 50 mM HEPES buffer followed by cold Dulbecco’s PBS (DPBS) before overlaying an ECM mixture of 0.05 mg/mL collagen I and 0.1 mg/mL fibronectin in DPBS. Devices were kept at 4°C in a humidified, sealed plate for 12-48 hr. On the day of EC seeding, devices were moved to 37°C and incubated for 1 hr to gel ECM. Excess ECM was then aspirated, channels were rinsed and filled with complete EC media, and devices were kept at 37°C until seeding. ECs were seeded at 4×10^6^ cells/mL (C166) or 5-7×10^6^ cells/mL (HUVEC) for mouse and human models, respectively. Cells were washed 2-4 hr post-seeding by adding media on top of inlet ports, and devices were cultured overnight to allow ECs to acclimate. On the following day, murine SVF or human RMFs were embedded in phenol red-free Matrigel (Corning) at a final concentration of 5×10^5^ cells/mL in 10 mg/mL Matrigel and seeded into the bottom channel of devices. Bottom channel inlet and outlet ports were then sealed with male mini luer plugs (Sigma) to prevent evaporation and maintain pressure across membrane during perfusion. Devices were maintained for one day under static conditions before connecting to perfusion.

### Perfusion Culture

NE-1600 or NE-1800 syringe pumps (New Era) were used for all experiments to provide consistent, controllable shear stress to the top channel of each MVoC. Syringes were filled with degassed EC media and secured in syringe pump. MVoCs were connected to the syringe pumps using 1/32 in tubing inserted into the inlet and outlet ports via 18G blunt tip needles. Prior to device connection, tubing was purged with media to expel any residual air bubbles formed during setup. A ramp-up program (**Table 1**) was then initiated, with MVoCs connected during the 10 µL/h phase and perfusion gradually increased to a continuous flowrate of 160 µL/h, corresponding to an estimated wall shear stress of 0.05 Pa. Wall shear stress was estimated using τ =6µQ/wh^2[42]^, where *µ* is dynamic viscosity (0.00093 Pa·s^[43]^), *Q* is volumetric flow rate, and *w* and *h* are channel width and height, respectively. To disconnect for routine maintenance, radiation, or endpoint analysis, syringe pumps were stopped and connections were carefully removed from device. To reconnect, pumps were purged and the ramp-up program was followed as with initial perfusion initiation. For longer-term experiments requiring replenishment of the syringes (>72 hr of perfusion), devices were disconnected, and the withdraw function of the syringe pump was used to refill syringes with fresh, degassed EC media.

**Table 1:**
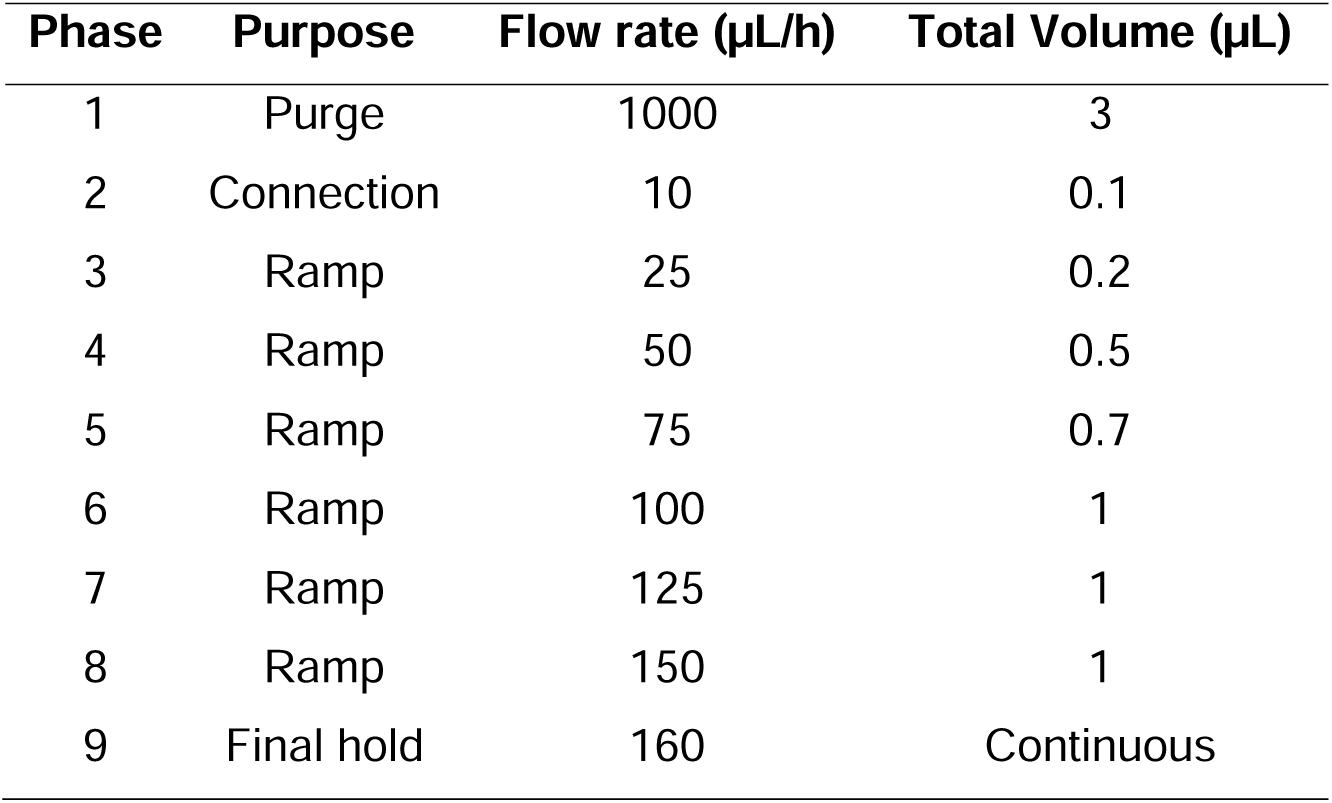
Ramp-up protocol for perfusion.

### Calcein-AM/Ethidium Homodimer-1 Staining

Cell viability in MVoCs was assessed using calcein-AM (Sigma) and ethidium homodimer (Invitrogen). Top channels of devices were washed by pipetting 150 µL of DPBS through top channels. 150 µL of DPBS containing 4 µM calcein-AM and 2 µM ethidium homodimer was then added to the top channel and devices were incubated in the dark at room temperature for 30 min. A tile scan of 20-22 fields of view spanning the straight portion of device channels was then captured using GFP (calcein-AM) and TXR (ethidium homodimer) filter cubes with a DMi8 inverted microscope (Leica). The number of live (calcein-AM) and dead (ethidium homodimer) cells was quantified using a threshold-based Fiji (ImageJ)^[44]^ macro and the viability of each device was calculated by dividing the number of live cells by total cells (live + dead) per image and averaging the viability per device.

### Immunofluorescence Staining

Top channels of devices were washed with 150 µL of DPBS. Top channels were fixed in methanol for 10 min at −20°C (ZO1, CD31) or in 10% neutral buffered formalin for 10 min at room temperature (γH2AX, p21). Devices were rinsed three times with DPBS. For γH2AX and p21 staining, devices were permeabilized with 0.1% Triton X-100 in DPBS for 5 min at room temperature. For all staining, channels were blocked for 1 hr at room temperature in DPBS with 1% bovine serum albumin, 22.5 mg/mL glycine, and 10% donkey (ZO1, CD31) or goat (γH2AX, p21) serum. The following primary antibodies were diluted in 1% BSA in DPBS and incubated overnight at 4°C: ZO1 (1:200; Novus NBP1-85047, lot no. 000046652), CD31 (1:100; R&D Systems AF3628, lot no. YZU0325062), γH2AX (1:1000; Invitrogen MA5-38225, lot no. ZL4579746B), anti-mouse p21 [E2R7A] (1:400; Cell Signaling 37543, lot no. 1), and anti-human p21 [12D1] (1:400; Cell Signaling 2947, lot no. 14). Devices were then rinsed three times with DPBS before incubating with secondary antibodies and 1.5µM Hoechst in 1% BSA in DPBS for 1 hr at room temperature. For F-Actin staining of RMFs and SVFs, pipette tips were filled with Phalloidin in 1% BSA in DPBS and gently placed into inlet and outlet ports if bottom channels for incubation with the secondary antibodies. Devices were then rinsed three times with DPBS and stored at 4°C until imaging. Images were captured using an inverted microscope (Leica) with GFP, TXR, DAPI, and Y5 filter cubes.

### Radiation

For γH2AX analysis, devices were exposed to a dose of 5, 10, or 20 Gy using a cesium source. For all other experiments, devices were exposed to a dose of 10 Gy using a cesium source. Sham controls (0 Gy) were transported with irradiated samples to ensure the same environmental conditions. Following exposure, top channels were rinsed with fresh media and devices were fixed (γH2AX) or reconnected as detailed above.

#### γH2AX analysis

Devices were fixed 30 min post-RT and stained for γH2AX as detailed above. For each device, γH2AX fluorescence was quantified from ten fields of view acquired at 64x magnification. Nuclear regions of interest (ROIs) were identified from the corresponding DAPI channel using a threshold-based macro in Fiji (ImageJ). The resulting ROIs were overlaid onto the corresponding γH2AX channel images and the maximum γH2AX intensity within each nucleus was quantified. For final analysis, 50 nuclei per device were selected using a random number generator. γH2AX intensity was normalized to the corresponding 0 Gy control within each experimental run to account for differences in baseline fluorescence intensity between independent experiments.

### FITC-dextran Permeability Assay

At each time point post-RT, top channels were rinsed with DPBS and 1 mg/mL of 40 kDa FITC-dextran (Chondrex) was added to each channel. Devices were incubated under static conditions for 30 min at 37°C in a humidified stage-top incubator (Oko-Lab). Fluorescence accumulation within the bottom channel gel was assessed via fluorescence microscopy at 0, 10, 20, and 30 min after adding the dextran. At each timepoint, a 10-slice z-stack spanning 10 µm was acquired, with field of view maintained consistently for each device throughout the assay.

For quantifying change in fluorescence intensity over time, z-stacks were converted to average-intensity projections and the mean fluorescence intensity was measured within a consistent region across all timepoints for each device. To correct for global changes in fluorescence intensity during image acquisition, ROI fluorescence was divided by the mean fluorescence intensity of the entire image at each timepoint (*I_ROI_*/*I_whole_*). The corrected intensity at each timepoint was then normalized to its corresponding baseline value (t=0) as (*I_t_-I_0_*)/I_0_. Permeability index (min^-1^) was calculated independently for each device as the slope of normalized fluorescence versus time during the initial linear diffusion interval (10-30 min).

*Senescence-associated beta-galactosidase activity (SA-*β*-gal) assay*

SA-β-gal activity was measured using the Senescence beta-Galactosidase Staining Kit (Cell Signaling) following the manufacturer’s protocol. Briefly, devices were fixed at room temperature for 10 min in 1X fixative (Cell Signaling) followed by three DPBS rinses before incubating in X-gal staining solution (pH 6.0) at 37°C for 24 hr. Representative brightfield images were acquired via a DMi8 inverted microscope (Leica).

### Conditioned Media Analysis

Effluent was collected from the outlet of murine MVoC top channels at various timepoints following RT to 0 or 10 Gy, which were maintained in DMEM supplemented with 2% FBS. Device effluent was collected over defined intervals post-RT as follows: 0-2 day effluent for 2 day samples and 5-7 day effluent for 7 day samples. Effluent was filtered using 0.22 µm PES syringe filters and aliquots were stored at −80°C until analysis. Cytokine profiling was performed by Eve Technologies Corporation (Calgary, AB, Canada) using a mouse 14-plex kit. Table S1 lists all cytokines analyzed. Of the cytokines in the multiplex panel, interleukin-6 (IL-6) and Chemokine C-C motif ligand 2 (CCL2) produced quantifiable signals suitable for comparison across experimental conditions and were included in subsequent analyses. The remaining analytes were below the assay detection range or were detected at concentrations comparable to basal media and were therefore excluded from quantitative analysis.

### Neutrophil Adhesion Assay

Neutrophil adhesion to endothelial monolayers was assessed under continuous perfusion at various timepoints post-RT. At each timepoint, neutrophils were labeled with 1 µM CellTracker Deep Red (Invitrogen) in serum-free DMEM for 20 min at 37°C. Cells were subsequently washed and resuspended in degassed DMEM/2% FBS for murine experiments or complete EGM-2 for human experiments. Differentiated MPRO and HL-60 cells were resuspended to concentrations of 2×10^6^ and 5×10^5^ cells/mL, respectively, and loaded into fresh syringes and tubing. Neutrophils were then perfused through the top channel of MVoCs for 30 min following the perfusion start-up protocol as detailed above. Following perfusion, adhesion to endothelial monolayers was imaged by acquiring a tile scan of 20-22 fields of view spanning the straight portion of each channel at 20x magnification using a Y5 filter cube. Adherent neutrophils in each field of view were quantified using a threshold-based macro in Fiji (ImageJ). 5 fields of view per device were randomly selected and included in the final quantitative analysis. The mean number of adherent neutrophils across the 5 fields was calculated for each device and used for subsequent statistical analysis and data visualization.

### Tumor Cell Adhesion Assay

Tumor cell adhesion to endothelial monolayers post-RT and following neutrophil adhesion was assessed under continuous perfusion. Immediately following the neutrophil adhesion assay, 4T1-dTom cells or CellTracker Green CMFDA-labeled (10 µM) MDA-MB-231 cells were collected and resuspended to a concentration of 5×10^5^ cells/mL in DMEM/2% FBS or complete EGM-2, respectively. Tumor cells were loaded into fresh syringes and tubing and perfused through MVoCs for 20 min following the perfusion start-up protocol as detailed above. Tumor cell adhesion was subsequently imaged and quantified as in neutrophil adhesion assays.

### Quantification and Statistical Analysis

All plots were made using GraphPad Prism. Imaging analysis was performed using Fiji (ImageJ). Statistical significance was determined via GraphPad Prism. Relevant statistical tests are indicated in figure legends. All experiments include at least n=3 independent devices collected from at least two independent experiments unless otherwise noted.

## Results

### Microfluidic device design and fabrication

During metastasis, circulating tumor and immune cells are recruited to the pre-metastatic niche where they adhere to the endothelium of microvasculature before extravasating into the surrounding tissue^[9,10]^. In breast tissue, this microvasculature is embedded within a fibrous stroma surrounding the mammary epithelial ducts and lobules, serving as the primary interface for cell trafficking and signaling between the circulation and mammary gland^[45,46]^ (**Figure 1A, B**). Because post-capillary venules are the principal sites of leukocyte and tumor cell extravasation^[47–49]^, we designed a microfluidic device to recapitulate this vascular-stromal interface. The device employs a stacked-channel geometry, which is a widely adopted microfluidic design for modeling tissue barriers^[50–53]^ that we adapted to recreate the mammary gland vessel microenvironment. In this model, ECs cultured under physiological shear stress in the upper channel are separated by a porous membrane from mammary fibroblasts embedded within an ECM hydrogel in the bottom channel (**Figure 1C**), replicating the vascular-fibrous stromal interface through which cells extravasate into the mammary gland.

**Figure 1.**
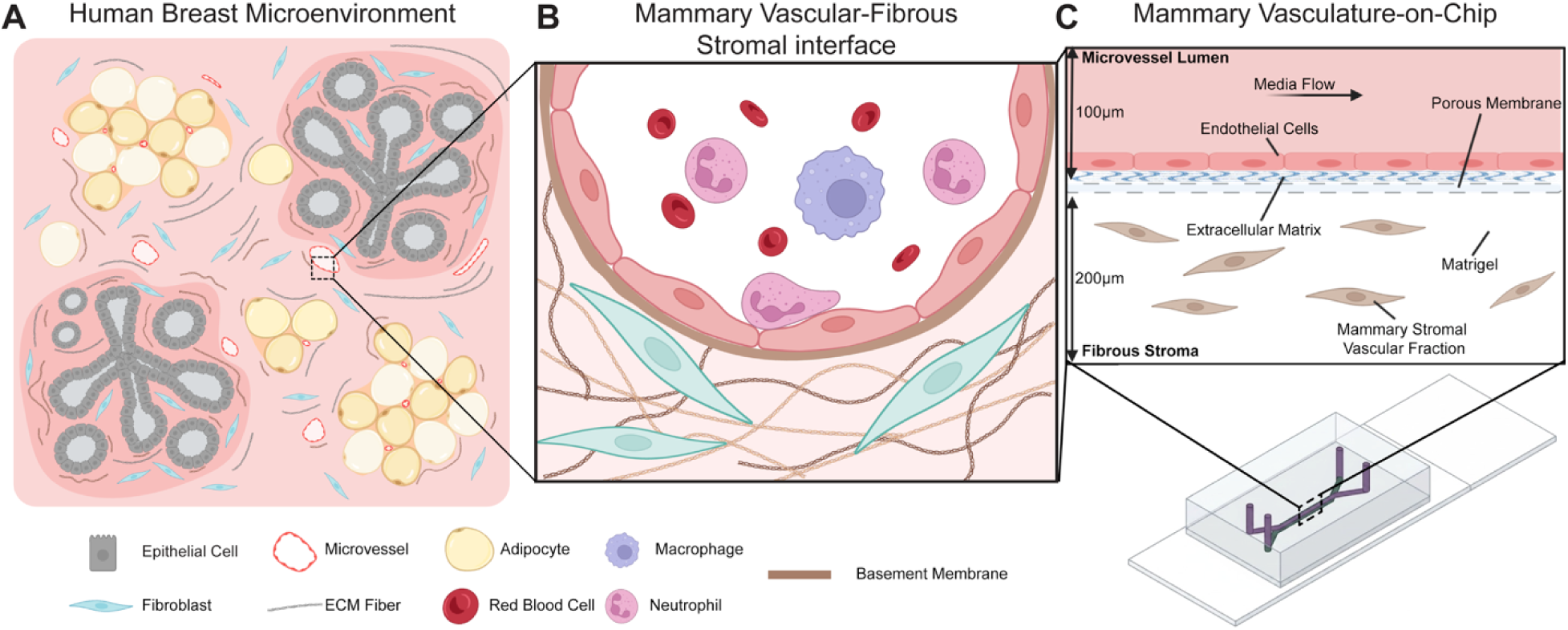
Mammary vasculature-on-chip (MVoC) recapitulates vascular-stromal interface of the breast. Schematics of the human breast microenvironment **(A)**, interface of a microvessel and the surrounding fibrous stroma **(B)**, and biomimetic vascular-stromal interface in the MVoC model **(C)**.

Our devices consisted of three layers: two PDMS slabs containing a 600 µm wide microchannel and a porous, PET membrane (**Figure 2A-B**). We used soft lithography to cast PDMS onto separate molds for the top and bottom channel features. The method of fabrication for these molds differed to create device layers with optimal features for each compartment. For the top channel, we 3D-printed features using a biocompatible resin, yielding molds with an average feature height of 106.7 ± 3.6µm (**Figure 2C-D**). We required a method that would yield precise and accurate feature heights in the top channel to minimize variation in shear stress across devices made from different features within a mold or using separate molds. Our 3D-printed molds each contained three features and yielded less than a 10% difference in shear stress across channels fabricated using different molds. For the bottom channel feature molds, we found that the optical clarity of our biocompatible resin was not high enough to clearly and consistently visualize cells in the devices. This agrees with previous reports of inherent surface roughness and subsequential low optical clarity generated by the additive manufacturing of 3D-printed molds^[54,55]^. We therefore used photolithography to fabricate SU-8 master molds with an average feature height of 183.6 ± 8.5 µm (**Figure 2D-E**). While these molds had a broader range of feature heights, each feature remained within 6% of the average height and did not impact cell behavior due to the static culture conditions. To ensure irreversible bonding between the top and bottom channel PDMS layers and the porous membrane, we treated the membrane with an amine-PDMS linker to functionalize the surface and make it compatible for irreversible bonding with PDMS^[56]^. We then used O_2_ plasma to successively bond each layer together before adhering the final device onto a microscope slide to improve stability of the PDMS during handling and allow for ease of imaging (**Figure 2F-G**).

**Figure 2.**
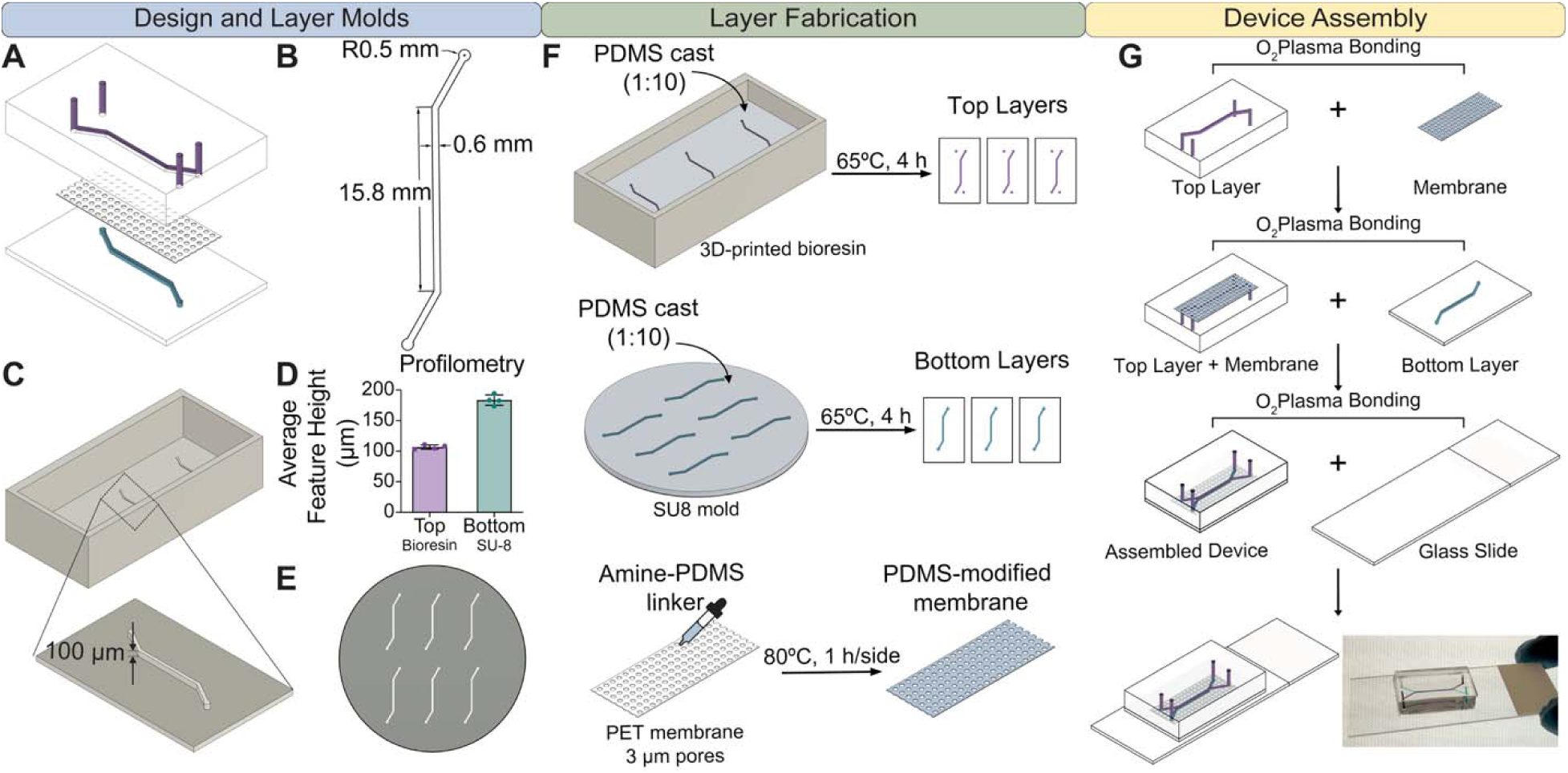
Microfluidic device design and fabrication. **A.** Exploded view of device, including top layer, porous membrane, and bottom layers. **B.** Channel dimensions for both top and bottom layers. **C.** Rendering of 3D-printed mold fabricated from biocompatible resin. **D.** Average feature heights across top and bottom layer molds measured via profilometry. Each point represents one feature. **E.** Rendering of SU-8 master for bottom layer channels. **F.** Layer fabrication steps. PDMS was cast at a 1:10 ratio of curing agent to base into top and bottom layer molds and cured. Top layers were cut from cured PDMS and hole punched, and bottom layers were cut and stored. Porous membranes were treated with amine-PDMS linker and stored. **G.** Schematic of device assembly steps. Layers and porous membrane were assembled successively and bonded to glass slides using O_2_ plasma.

### MVoCs model the vascular-stromal interface under shear stress

Endothelial behavior and circulating cell adhesion are highly regulated by fluid shear stress, underscoring the importance of establishing a physiologically relevant model of the vascular-stromal interface to study immune and tumor cell adhesion during metastasis^[57–60]^. We cultured ECs in the top channel of MVoCs under perfusion to achieve a shear stress of 0.05 Pa, which is the amount of shear experienced by ECs in the post-capillary venule during peak neutrophil adhesion^[61–63]^, making it ideal for studying immune and tumor cell dynamics. This shear stress was well-tolerated by both murine and human ECs, as they maintained at least 97% viability over 8 days under perfusion, with 99% viability by day 8 (**Figure 3A, B**). Both murine and human ECs maintained CD31 expression and formed tight junctions when cultured under shear stress in our devices (**Figure 3C**).

**Figure 3.**
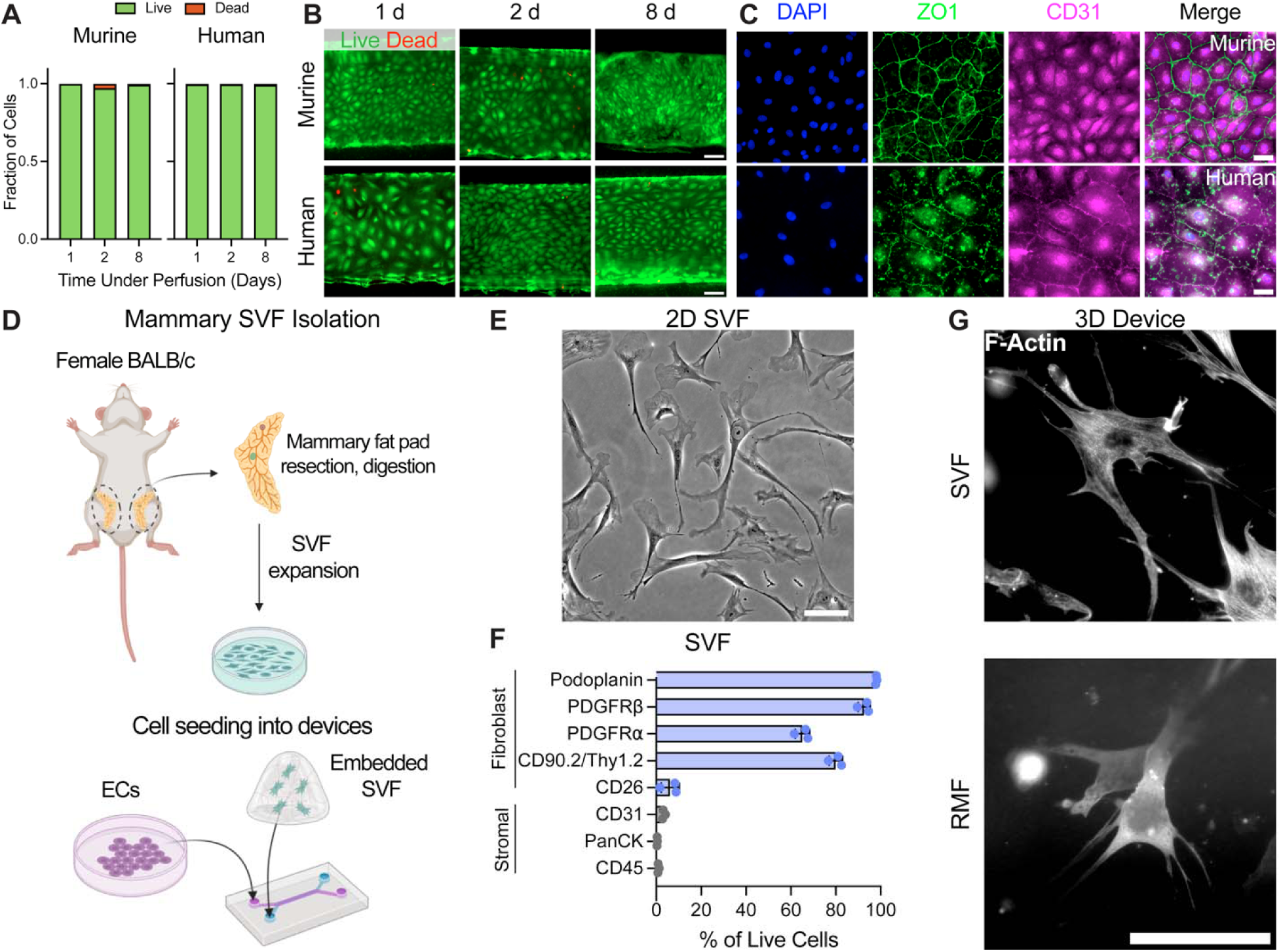
Cell culture in MVoCs. **A.** Quantification of murine and human EC viability under perfusion in MVoCs via Live/Dead (calcein-AM/ethidium homodimer-1) staining with representative images under perfusion in (**B)**. Scale bars are 100 µm. **C.** Representative images of tight junction (ZO1) and an endothelial cell marker (CD31) in MVoCs under perfusion. Scale bars are 50 µm. **D.** Schematic of stromal vascular fraction (SVF) isolation from murine mammary tissue. Mammary fat pads were harvested from female BALB/c mice, digested, and plated for expansion. Cells were embedded into Matrigel and seeded into the bottom channel of MVoCs. **E.** Representative phase contrast image of SVF in 2D culture prior to seeding into MVoCs. Scale bar is 100 µm. **F.** Flow cytometry characterization of SVF after expansion and before seeding in MVoCs. n=3 mice. **G.** Representative images of F-Actin staining of SVF and RMFs embedded in Matrigel within MVoCs. Scale bar is 100 µm.

Fibroblasts are a highly heterogenous population within breast tissue that exert a range of functions, supporting development, remodeling, and homeostasis of the mammary gland in both humans and mice^[64–66]^. To create a breast model and understand the contribution of mammary-specific cells in the vascular response to RT, we isolated the stromal vascular fraction (SVF) from mammary fat pads of naïve BALB/c mice (**Figure 3D**). At the time of embedding into the devices, the SVF appeared homogeneous with an elongated, spindle-like morphology (**Figure 3E**). SVF cells expressed general fibroblast proteins, including platelet-derived growth factor receptor alpha and beta (PDGFR, PDGFR), podoplanin, and CD90.2/Thy1.2, with few cells expressing CD26 and negligible expression of immune, epithelial, and EC markers (**Figure 3E; Supplemental Figure S1A, B**). This suggests that the SVF in MVoCs was comprised primarily of fibroblast-like cells as we previously reported^[16]^. To model the human breast microenvironment, we used hTERT-immortalized reduction mammoplasty fibroblasts (RMFs)^[41]^. These cells also displayed a spindle-like morphology and expressed negligible E-cadherin with high levels of vimentin (**Supplemental Figure S1C-E**). RMFs also exhibited baseline expression of fibroblast activation protein (**Supplemental Figure S1F**), thereby providing a model of fibroblast-like cells for our human MVoC. When cultured in the 3D ECM gel within the bottom channel of MVoCs, both SVF cells and RMFs exhibited visible actin protrusions (**Figure 3G**). Together, this establishes human and murine MVoCs that contain a viable, tight junction-forming endothelium cultured under physiological shear stress in indirect co-culture with a mammary-specific fibrous stroma.

### Modeling the mammary vascular response to RT

Clinical data strongly supports that RT prevents recurrence in a majority of breast cancer patients^[1]^. RT does not spare normal tissue, however, and in patients that do experience recurrence, a pre-metastatic niche may form in this radiation-damaged mammary tissue to drive the infiltration of immune and tumor cells^[4,16,67]^. We therefore wanted to understand how the ECs in MVoCs respond to RT when exposed to shear and co-cultured with mammary fibroblasts. After overnight perfusion, we irradiated MVoCs and continued to culture them under shear stress for both short- and longer-term periods, replicating acute and delayed responses to RT (**Figure 4A**). Both murine and human ECs exhibited a dose-dependent increase in DNA double-strand breaks when exposed to doses from 0 to 20 Gy, as evidenced by the increasing intensity of γH2AX foci per nucleus at 30 min post-RT (**Figure 4B, C**). While 20 Gy is consistent with both intra- and post-operative RT in breast cancer patients and is used for *in vivo* studies of pre-metastatic niche formation in the normal mammary tissue^[4,68,69]^, we used 10 Gy for our remaining studies as this is the maximum dose that we are able to observe molecular changes in *in vitro* 2D models without significant cell death^[4,70]^. At this dose, ECs exhibited minimal shifts in viability up to 7 days post-RT (**Figure 4D, E**). Barrier integrity is a critical function of the endothelium and can impact cell extravasation^[71]^, especially after treatment with RT or other therapeutics. We analyzed changes in endothelial monolayer permeability post-RT by measuring the rate of FITC-dextran diffusion from the vascular channel into the underlying stromal compartment. Endothelial permeability began to increase as early as 6 hr post-RT and peaked with a 4-fold increase in permeability index at 1 day before returning to baseline by 3 days (**Figure 4F**). This initial damage and recovery was also exhibited through visible decreases in ZO1 expression in both murine and human monolayers, which appeared to be more segmented and reduced in intensity at 1 day post-RT in 10 Gy compared to 0 Gy devices (**Figure 4G, H**). Taken together, both our murine and human MVoC models demonstrate that endothelial monolayers in co-culture with mammary fibroblasts exhibit an acute response consistent with a typical initial spike in DNA damage and permeability followed by recovery of the monolayer^[11,72]^, highlighting the utility of using these models to understand the mammary vascular response to RT.

**Figure 4.**
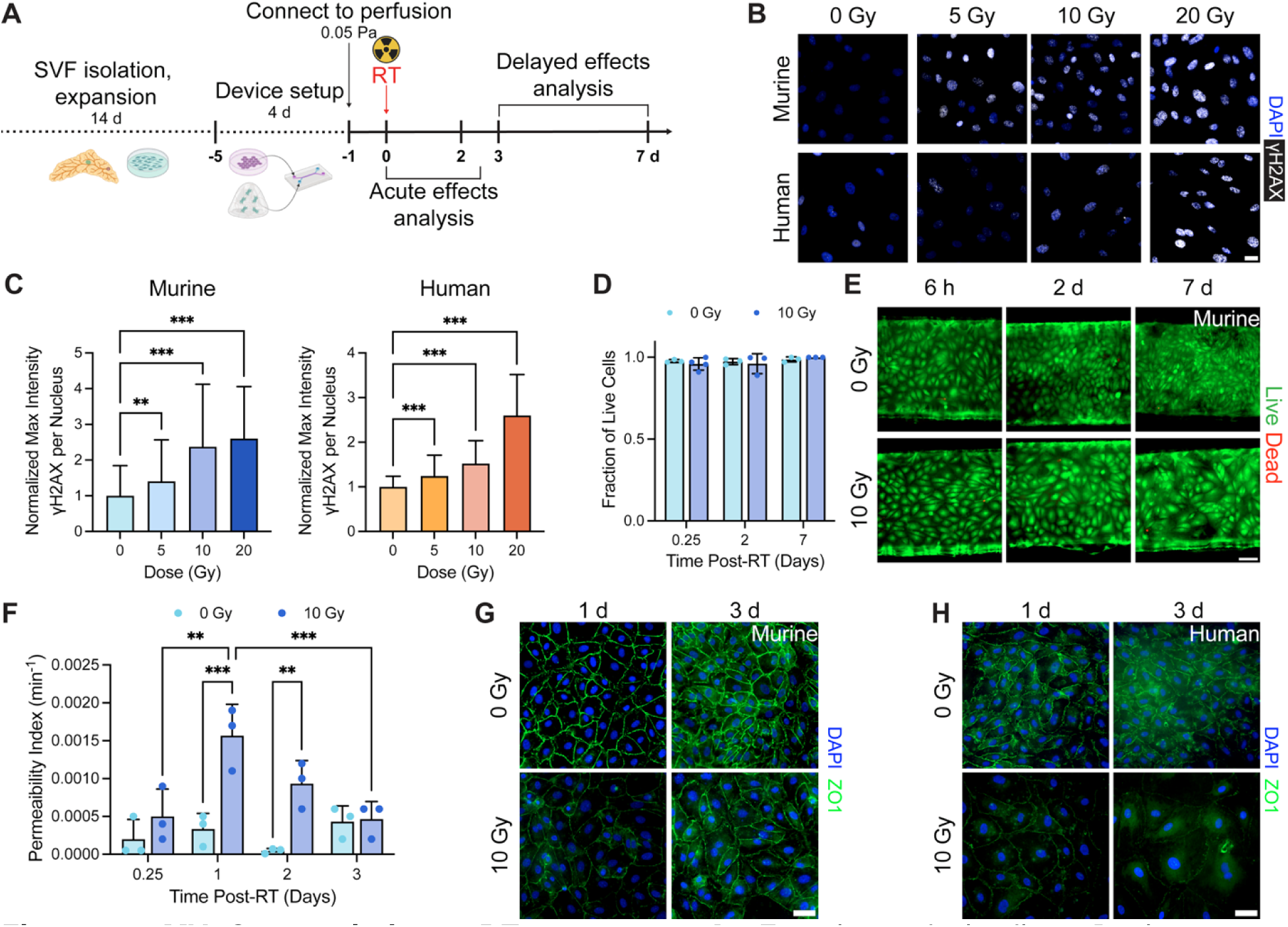
MVoC recapitulates RT response. **A.** Experimental timeline. Devices were connected to perfusion overnight prior to exposure to radiation. Acute (1-2 d) and delayed (3-7 d) effects of radiation on the mammary chips were analyzed. Representative images **(B)** and quantification of murine and human γH2AX staining **(C)**. Scale bar is 25 µm. n=3 independent chips per condition with 50 nuclei analyzed per chip. **p<0.01 and ***p<0.001 via Kruskal-Wallis test with Dunn’s test for multiple comparisons. **D.** Quantification of murine EC viability post-RT. n=3 independent chips per condition. **E.** Representative images of Live/Dead staining at 6 h, 2 d, and 7 d post-RT in murine devices. Scale bar is 100 µm. **F.** Permeability index of murine EC monolayers post-RT measured via 40 kDa FITC-dextran diffusion into the bottom channel. n=3 independent mammary chips per condition. **p<0.01 and ***p<0.001 via two-way ANOVA with Tukey’s test for multiple comparisons. Representative images of ZO1 expression in murine **(G)** and human **(H)** EC monolayers at 1 and 3 d post-RT. Scale bar is 50 µm. All data show mean ± SD.

While the endothelium can repair itself physically and restore its barrier integrity post-RT, it is also known that senescence is a critical molecular response of normal tissue to radiation exposure that can modulate the microenvironment and have significant impacts on inflammation^[6,11,73,74]^. We therefore sought to characterize the senescence response of our devices post-RT. Within 2 days, both murine and human ECs exhibited hypertrophy as evident by an increase in cell area (**Supplemental Figure S2A-B**). Interestingly, 10 Gy human ECs maintained a 1.6-fold increase in cell size compared to 0 Gy at both 2 and 7 days post-RT, while 10 Gy murine ECs exhibited only a slight increase in size at 2 days (1.1-fold) followed by a 2.6-fold increase by 7 days post-RT. We also observed increased β-galactosidase (β-gal) activity and cell cycle arrest in both ECs and mammary fibroblasts post-RT (**Figure 5A-D; Supplemental Figure S2C-D**). Low-level β-gal activity was present in 0 Gy devices, likely due to the confluent monolayers^[75]^. However, differences between 0 and 10 Gy became more apparent over time as the intensity of the β-gal staining increased and became more concentrated rather than diffuse. Irradiated ECs in human MVoCs exhibited stronger β-gal activity earlier than those in our murine model, with overall more concentrated staining observed across irradiated conditions.

**Figure 5.**
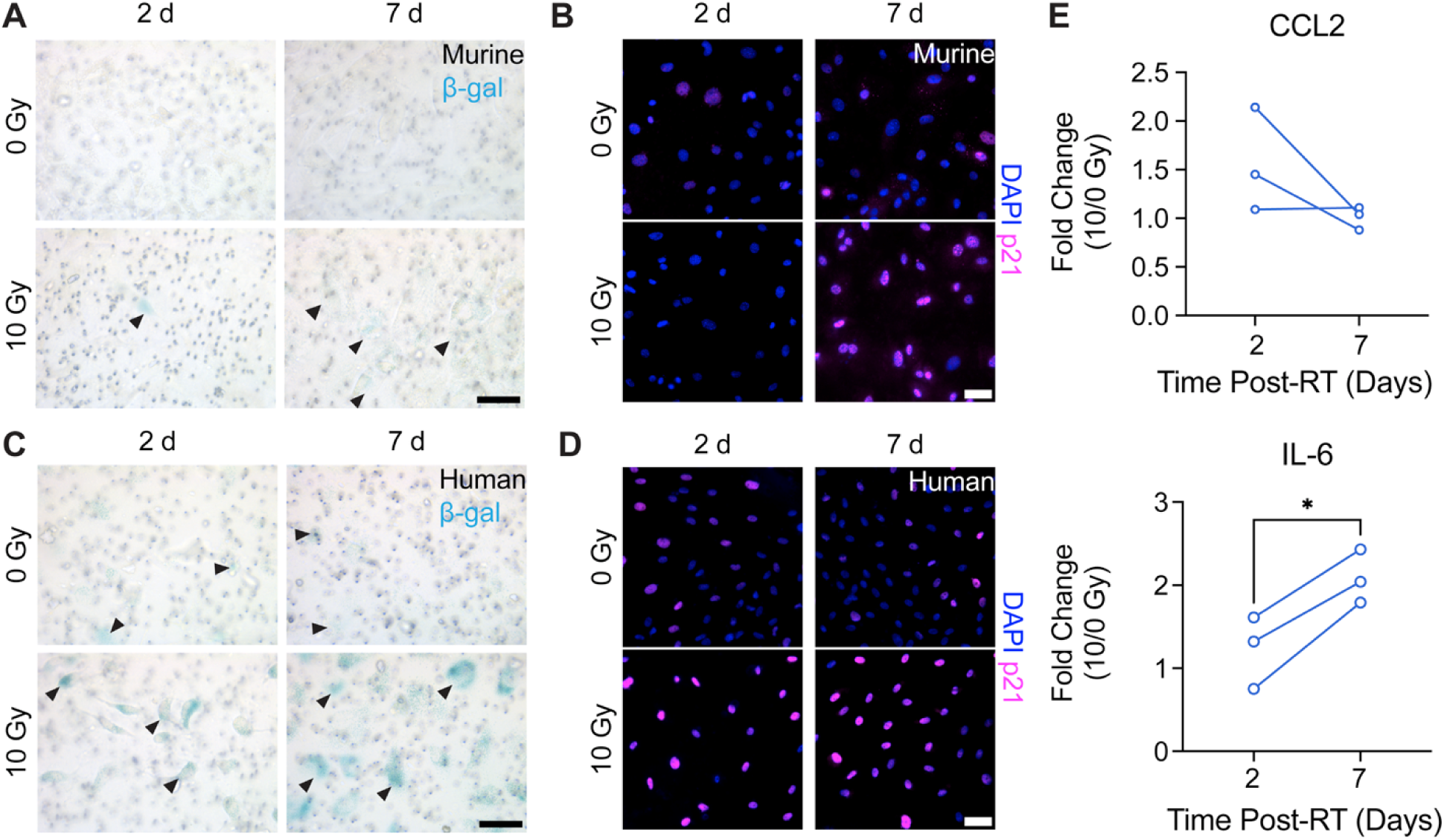
Murine and human MVoCs exhibit senescent-like phenotype post-RT. **A.** Representative brightfield images of β-galactosidase (β-gal) staining of murine EC monolayers post-RT. Arrows indicate positive β-gal signal. Scale bar is 100 µm **B.** Representative images of p21 staining of the murine endothelium post-RT. Scale bar is 50 µm. **C.** Representative brightfield images of β-gal staining of human EC monolayers post-RT. Arrows indicate positive β-gal signal. Scale bar is 100 µm **D.** Representative images of p21 staining of the human EC post-RT. Scale bar is 50 µm. **E.** Murine cytokine analysis of conditioned media from the vascular compartment of MVoCs post-RT. n=3 independent chips per condition. *p<0.05 via paired t-test.

Secretion of pro-inflammatory factors as part of a senescence-associated secretory phenotype (SASP) is also characteristic of both ECs and fibroblasts post-RT^[6,74]^. Consistent with this response, irradiated murine MVoCs exhibited increased secretion of the SASP-associated factors CCL2 and IL-6 relative to time-matched 0 Gy controls (**Figure 5E**). CCL2 secretion was increased at 2 days post-RT before a reduction back to control secretion levels by day 7. In contrast, the RT-associated increase in IL-6 secretion was most pronounced at 7 days, suggesting temporally distinct changes in the MVoC secretory response post-RT. Together, these findings demonstrate the development of an RT-induced senescent phenotype at the vascular-stromal interface characterized by cellular hypertrophy, cell cycle arrest, β-gal activity, and time-dependent changes in secretion of SASP-associated inflammatory factors.

### Post-RT neutrophil adhesion sustains endothelial damage and increases tumor cell adhesion

Neutrophils play a critical role in the acute wound healing response but have also been implicated in the longer-term process of promoting tissue damage that facilitates a pre-metastatic niche^[7,76,77]^. As proof-of-concept for using our mammary vascular chip to study immune and tumor cell interactions with the endothelium and stroma post-RT, we sought to understand neutrophil adhesion dynamics on the irradiated ECs and how neutrophils may impact downstream processes like barrier recovery and tumor cell infiltration. We perfused neutrophils through devices for 30 min up to 3 days after RT to determine the peak of neutrophil adhesion (**Figure 6A**). We found that this length of perfusion allowed analysis of adhered neutrophils before they detached (**Supplemental Figure S3**). Starting at 6 hr, irradiated devices showed increased neutrophil adhesion compared to controls in murine MVoCs (**Figure 6B**). This adhesion peaked at 2 days post-RT before returning to baseline at 3 days, at which point we saw no difference in adhesion between control and irradiated MVoCs. Interestingly, some neutrophils that adhered to irradiated monolayers after 2 days appeared to be elongated and spread out (**Figure 6C**), suggesting activation and potential NETosis^[78]^, though additional characterization is required for confirmation. In our human model, we also observed increased neutrophil adhesion to endothelial monolayers after 2 days (**Figure 6D**).

**Figure 6.**
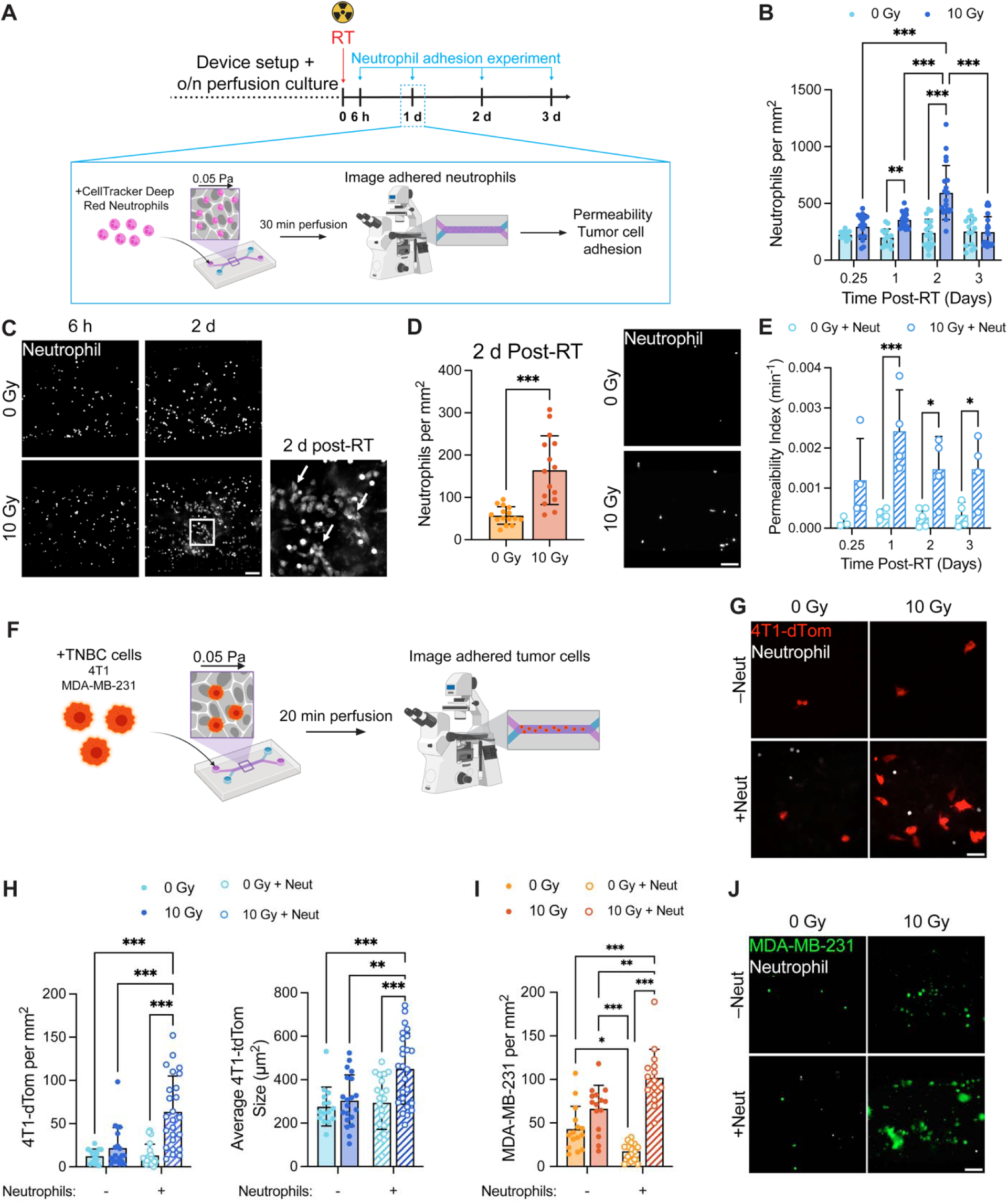
RT-mediated neutrophil increases tumor cell attachment to the endothelium. **A.** Experimental timeline. Devices were irradiated and neutrophils were perfused for 30 min at 6 h and 1, 2, and 3 d post-RT. Devices were imaged to quantify neutrophil adhesion before proceeding to permeability and tumor cell adhesion assays. Quantification **(B)** and representative images of neutrophil (dMPRO) adhesion **(C)** in murine MVoCs. Data show mean ± SD, where each point is one representative image with 10 images and n=3-4 independent chips per condition. Scale bars are 100 µm. **p<0.01 and ***p<0.001 via two-way ANOVA with Sidak’s test for multiple comparisons. **D.** Permeability index of murine EC monolayers post-RT and post-neutrophil adhesion measured via 40 kDa FITC-dextran diffusion into the bottom channel. n=3-5 independent murine MVoCs per condition. *p<0.05 and ***p<0.001 via two-way ANOVA with Tukey’s test for multiple comparisons. **E.** Quantification and representative images of neutrophil (dHL-60) adhesion in human MVoCs at 2 d post-RT. Each point represents one representative image with 10 images and n=3 independent chips per condition. ***p<0.001 via Mann-Whitney test. Scale bar is 100 µm. **F.** Experimental timeline for tumor cell adhesion assay. Following neutrophil perfusion, 4T1-dTom or CellTracker Green-stained MDA-MB-231 cells were perfused for 20 min through MVoCs before imaging to quantify the number of adhered tumor cells to the endothelium. **G.** Representative images of tumor cell adhesion in murine MVoCs. Scale bar is 50µm. **H.** Quantification of 4T1-dTom (red) adhesion and size in murine MVoCs. Each point is one representative image with 10 images and n=3-4 independent chips per condition. Quantification **(I)** and representative images **(J)** of MDA-MB-231 (green) adhesion in human MVoCs. Each point is one representative image with 10 images and n=3 independent chips per condition. *p<0.05, **p<0.01, and ***p<0.001 via two-way ANOVA with Tukey’s test for multiple comparisons. Scale bar is 100 µm.

While neutrophils initiate wound healing to repair tissue, they have also been shown to prolong damage and induce immunosuppressive, pro-tumor signaling when pathologically activated^[7,76]^. This activation can create a feedback loop between continuously damaged vasculature and epithelium, neutrophil activation, and incomplete resolution of tissue damage. We therefore sought to understand the recovery dynamics of the endothelial barrier when exposed to neutrophils. Neutrophils significantly increased permeability in irradiated compared to unirradiated murine MVoCs from 1 to 3 days post-RT (**Figure 6E**) in contrast to the peak in permeability 1 day and subsequent recovery that we observed by 3 days with RT alone (**Figure 4F**). This sustained increase in permeability from 6 hr to 3 days post-RT suggests that neutrophils may delay recovery of the endothelium.

Neutrophils and other immune cells have been shown to influence tumor cell seeding by increasing survival in circulation, aiding adhesion to the endothelium, and assisting with invasion into the metastatic niche^[79]^. Knowing that neutrophil adhesion peaked at 2 days post-RT in our system, we sought to understand how tumor cells would adhere to the endothelium when pre-conditioned with neutrophils. Following neutrophil perfusion and adhesion, we perfused 4T1-dTom or MDA-MB-231 TNBC cells through murine and human devices, respectively, for 20 min, which we observed to be the plateau of tumor cell adhesion post-RT and post-neutrophil adhesion before declining (**Figure 6F; Supplemental Figure S4; Supplemental Video S1**). Irradiated MVoCs that were exposed to neutrophils showed significantly more tumor cell adhesion compared to unirradiated and MVoCs without neutrophils in both murine and human models (**Figure 6G-J**). Furthermore, tumor cells that adhered to irradiated, neutrophil-conditioned murine ECs were significantly larger on average (**Figure 6H**), suggesting that the combination of RT and neutrophils not only enables more tumor cell adhesion but also increases the speed at which tumor cells spread on the vascular barrier. Of note, neutrophils appeared to exhibit a protective effect in unirradiated human MVoCs, as we observed a significant reduction in tumor cell adhesion to the ECs of 0 Gy human MVoCs pre-conditioned with neutrophils compared to those with tumor cells alone (**Figure 6I**). Taken together, these studies highlight the pro-metastatic role that neutrophils may play in irradiated breast tissue.

## Discussion

With the incidence of breast cancer rising annually, development of new treatments and improved therapeutic protocols is imperative. Nearly one in three patients will experience recurrence, which may be caused by CTCs that shed from the primary tumor before and during initial treatment before infiltrating the resected primary tumor bed of mammary tissue or colonizing distant organs (e.g., lung, liver, bone)^[4,7,8,76]^. To develop effective treatments that prevent recurrence, it is crucial that we understand how pre-metastatic niches form in both local and distant organs. Because up to 30% of TNBC patients will experience locoregional failure post-RT^[2,4]^, we developed a MVoC model of the mammary vascular-stromal interface to understand how RT alters the vascular niche to favor CTC seeding. We found that RT causes transient barrier injury but a persistent phenotypic shift, potentially toward a SASP. Moreover, neutrophil conditioning and adhesion to MVoC EC monolayers prolonged barrier injury and increased tumor cell adhesion to irradiated MVoCs, suggesting that immune cell recruitment post-RT may modify the endothelium in a way that favors CTC capture and adhesion.

The endothelium is a major regulator of organ response to RT that undergoes both acute changes and chronic adaptations as a protective mechanism^[35,73,80]^. We found that human and murine MVoCs exhibited an acute response consistent with typical radiation exposure: increased γH2AX foci, barrier permeability, and leukocyte adhesion within the first 2 days. Additional changes were visible after 2 days, most notably in human MVoCs, including cellular hypertrophy, cell cycle arrest, and increased β-gal activity. This molecular shift toward a senescent phenotype, which persisted in the human MVoCs and became increasingly visible in the murine model by 7 days post-RT, suggests that longer-term adaptations may occur at the mammary vascular-stromal interface despite recovery from the physical effects of RT. This effect has been observed in other organ models both *in vivo* and *in vitro*^[38,81,82]^, where ECs remain viable but show signs of altered molecular signaling in the days to weeks post-RT.

However, this has not been extensively studied in mammary tissue models. Senescence and SASPs have been linked to chronic inflammation of the vascular barrier, particularly when innate immune cells are continuously recruited to and activated by the initial site of damage. In line with this phenotype, we observed increased IL-6 secretion from irradiated murine MVoCs that was most pronounced by 7 days post-RT. IL-6 is a canonical SASP-associated factor with established roles in immune regulation and stromal remodeling^[16,83]^. This phenotype may therefore be critical for pre-metastatic niche formation in irradiated mammary tissue, where we have previously observed chronic innate immune infiltration that precedes tumor cell seeding^[4,16]^. Further characterization of the potential EC SASP and additional molecular changes (e.g., Notch signaling, adhesion molecule regulation^[84]^) in MVoCs could identify potential targets to halt chronic endothelial inflammation and shift any pro-tumor adaptations that may drive pre-metastatic niche formation in the vascular niche.

Circulating immune cells must interact with the endothelium prior to extravasating into the tissue stroma. Alterations to EC phenotype and subsequent changes to their adhesion molecule expression, secretome, and molecular signaling will therefore affect immune cell recruitment, extravasation, and function^[85–87]^. We observed increased neutrophil adhesion to MVoC ECs as early as 6 hr post-RT, with a peak occurring at 2 days, which is consistent with the established acute window of leukocyte response and endothelial activation post-RT^[88–90]^. This coincided with increased CCL2 secretion from MVoCs between 0-2 days post-RT. While CCL2 is conventionally associated with monocyte recruitment, it also has a demonstrated role in neutrophil recruitment and activation during inflammation^[91,92]^. The coexistence of increased CCL2 secretion and maximal neutrophil adhesion at 2 days post-RT suggests that RT-induced changes to the endothelial secretome accompany acute immune cell recruitment. Moreover, we observed diffuse extracellular material and elongation of adhered neutrophils in irradiated MVoCs, which is suggestive of neutrophil extracellular trap (NET) release and neutrophil activation^[78]^. This demonstrates that neutrophils may become activated or shift their phenotype in response to the activated endothelium, though additional validation of NETosis and investigation into neutrophil function are needed for confirmation. Together, these findings suggest that the acute MVoC response to RT creates a transient, inflammatory vascular niche that facilitates neutrophil adhesion and may subsequently alter their function.

Pathologically activated neutrophils can cause persistent damage to the endothelium and surrounding tissue by releasing reactive oxygen species (ROS), NETs, and proteinases^[76,93,94]^. While we observed a peak in permeability and full recovery of endothelial barrier integrity by 3 days after RT alone, irradiated MVoCs that were also exposed to neutrophils showed an increase in permeability at every timepoint up to 3 days. This persistent permeability increase occurred only in MVoCs with neutrophil adhesion, suggesting that the neutrophils may be prolonging damage due to activation by the dysfunctional endothelium.

With evidence of a persistent shift in EC phenotype, our findings establish a potential positive feedback loop at the mammary vascular-stromal interface whereby the ECs are activated by RT and their initial activation and change in secretome serves to increase the adhesion of neutrophils. This may then cause inflammatory signaling and activation of the infiltrating neutrophils, which would release damaging molecules like ROS or enzymes that prolong the endothelial dysfunction and continue this signaling loop, ultimately forming an inflammatory, pre-metastatic niche in mammary tissue. The possibility of this pre-metastatic niche formation at the mammary vascular-stromal interface is further underscored by our finding that in both murine and human MVoCs, TNBC cells adhered significantly more to EC monolayers that had been both irradiated and pre-conditioned with neutrophil adhesion. Interestingly in the human model, neutrophils appeared to have a protective effect on unirradiated MVoCs, highlighting the likely dependency of neutrophil function on the state of the endothelium. Together, our findings demonstrate that interactions between irradiated ECs and neutrophils can alter subsequent tumor cell adhesion, supporting a key role for the vascular-stromal interface in development of a pre-metastatic niche post-RT.

Shear stress plays a major role in regulating the endothelial response to stress and their interaction with circulating cells^[57–60]^. Previous studies have used OoCs to study organ-specific RT responses in various organs like the liver, lung, and gut, employing shear and stretch to ensure physiologically-relevant cues for the tissue microenvironment^[35,38,53]^. Additional studies have also been performed to analyze the vascular response to RT^[95]^. While these studies are critical for laying the foundation for use of OoCs in radiobiology, there remains a gap in models with mammary-specific stromal microenvironments with a defined and easily tunable shear stress. Our MVoCs enable the study of interactions between RT-induced endothelial damage, circulating immune cells, and tumor cells, allowing the study of each component to be controlled independently over time. To our knowledge, this is the first model of normal mammary tissue that incorporates physiologic shear stress to study the effects of RT and how it may promote pre-metastatic niche formation. Fabricated using standard OoC protocols and commercially available materials, our MVoC aims to be technically and financially accessible for use in future radiobiological studies.

The modularity of our MVoCs also provides opportunity to improve the physiological complexity of the model and expand the stages of metastasis that can be investigated. Both our murine and human models do not include mammary-specific ECs, and our human model employs large vessel ECs. The addition of primary breast microvascular ECs from human donor tissue and isolation of ECs from murine mammary glands would advance the organ-specificity of our model and subsequently improve translation of findings. Despite this limitation, we observed persistent EC changes for at least 7 days post-RT. Given the ability of ECs to survive longer-term under shear stress than in static conditions, these studies could likely be extended to understand how chronic immune infiltration over a period of weeks impacts irradiated mammary tissue. Tumor cell behavior is also likely to be dependent on the persistent change in EC response. Expanding analysis of tumor cell behavior to include invasion and extravasation from the vascular compartment into the fibrous-stromal compartment would improve our understanding of this dependency and could be accomplished by expanding the pore size of MVoC membranes and extending the perfusion time of tumor cells beyond the 30 min timeframe used here to study adhesion. While the focus of this study was primarily on the EC response to RT, investigation into the fibroblast response would better elucidate the dynamic breast microenvironment post-RT and how it may enable a pre-metastatic niche. Furthermore, fibroblasts are heterogeneous within the mammary gland, and their response to stimuli can vary greatly depending on their basal expression of certain proteins^[64–66,96]^. Further characterization of both the SVF and RMFs would inform the direction of this RT response and ensure appropriate interpretation of these results. Together, these modifications would expand our MVoC from a model of early vascular recruitment and adhesion to one capable of interrogating long-term interactions that govern progression of the mammary pre-metastatic niche.

## Conclusion

Despite advancements in first-line therapies, there remain limited options for recurrence prevention in breast cancer. Our MVoC aims to replicate a key step of the metastatic cascade by modeling the vascular-stromal interface through which tumor cells must pass to colonize irradiated breast tissue. This work highlights the importance of studying interactions between immune and tumor cells, as their reciprocal effects are complex and dependent on their microenvironment. MVoCs can be used to probe molecular and cellular interactions in both murine and human platforms to reduce the reliance on animal models and improve the translational relevance of pre-clinical studies. Future incorporation of patient-derived tissues could further enable a personalized medicine approach by investigating interpatient variability and predicting RT response and susceptibility to recurrence. Ultimately, MVoCs and physiologically relevant models of irradiated mammary tissue will provide valuable pre-clinical platforms for identifying mechanisms of and therapeutic targets for recurrence.

## Author contributions

Conceptualization: S.E.M, M.R.; methodology: S.E.M., K.D., G.M.F., M.M.V.; investigation: S.E.M., S.N.L., K.D., G.M.F., M.P., M.M.V.; visualization: S.E.M.; supervision: M.R., L.M.B.; writing: S.E.M.; and review & editing: all authors.

## Acknowledgements

We thank Dr. Christina McGahan, Megan Dernberger, and the Vanderbilt Institute for Nanoscale Science and Engineering (VINSE) for their guidance in developing the microfluidic fabrication protocol and use of cleanroom equipment, Dr. Michael Freeman for irradiator use, Dr. Alissa Weaver for flow cytometer use, Dr. Charlotte Kuperwasser for providing immortalized reduction mammary fibroblasts, and Dr. Laura L. Bronsart for providing the dTomato-labeled 4T1 cells. The Vanderbilt Flow Cytometry Shared Resource administers the FlowJo Portal Group License used to analyze the flow cytometry data and is supported by the Vanderbilt Ingram Cancer Center (P30CA068485) and the Vanderbilt Digestive Disease Research Center (DK058404). This research was financially supported by the Vanderbilt Institute for Clinical and Translational Research Grant Number VR73788, a VINSE pilot award, National Institutes of Health Grant Numbers R00CA201304 (M.R.), F31CA298725 (S.E.M), and T32DK101003 (K.D.), and a Vanderbilt Scaling Success Award. Schematics were created using BioRender.

## References

[1] A. E. Dragun, J. Pan, D. Jain, B. Kruse, S. Rai, Int. J. Radiat. Oncol. 2009, 75, S216.

[2] A. J. Lowery, M. R. Kell, R. W. Glynn, M. J. Kerin, K. J. Sweeney, Breast Cancer Res. Treat. 2012, 133, 831.

[3] A. Afghahi, N. Purington, S. S. Han, M. Desai, E. Pierson, M. B. Mathur, T. Seto, C. A. Thompson, J. Rigdon, M. L. Telli, et al., Clin. Cancer Res. 2018, 24, 2851.

[4] M. Rafat, T. A. Aguilera, M. Vilalta, L. L. Bronsart, L. A. Soto, R. Von Eyben, M. A. Golla, Y. Ahrari, S. Melemenidis, A. Afghahi, et al., Cancer Res. 2018, 78, 4241.

[5] E. K. Rofstad, B. Mathiesen, K. Henriksen, K. Kindem, K. Galappathi, Cancer Res. 2005, 65, 2387.

[6] H. J. Hwang, Y. R. Lee, D. Kang, H. C. Lee, H. R. Seo, J. K. Ryu, Y. N. Kim, Y. G. Ko, H. J. Park, J. S. Lee, Cancer Lett. 2020, 490, 100.

[7] E. Nolan, V. L. Bridgeman, L. Ombrato, A. Karoutas, N. Rabas, C. A. N. Sewneth, M. Vasquez, F. S. Rodrigues, S. Horswell, P. Faull, et al., Nat. Cancer 2022, 3, 173.

[8] B. Ruiz-Fernández de Córdoba, H. Moreno, K. Valencia, N. Perurena, P. Ruedas, T. Walle, A. Pezonaga-Torres, J. Hinojosa, E. Guruceaga, A. Pineda-Lucena, et al., Cancer Discov. 2022, 12, 1356.

[9] H. Peinado, H. Zhang, I. R. Matei, B. Costa-Silva, A. Hoshino, G. Rodrigues, B. Psaila, R. N. Kaplan, J. F. Bromberg, Y. Kang, et al., Nat. Rev. Cancer 2017, 17, 302.

[10] D. X. Nguyen, P. D. Bos, J. Massagué, Nat. Rev. Cancer 2009, 9, 274.

[11] B. Baselet, P. Sonveaux, S. Baatout, A. Aerts, Cell. Mol. Life Sci. 2019, 76, 699.

[12] J. M. Brown, Int. J. Radiat. Oncol. Biol. Phys. 2020, 108, 734.

[13] M. De Palma, C. E. Lewis, Cancer Cell 2013, 23, 277.

[14] H. J. Park, R. J. Griffin, S. Hui, S. H. Levitt, C. W. Song, Radiat. Res. 2012, 177, 311.

[15] B. C. Hacker, J. D. Gomez, C. A. Silvera Batista, M. Rafat, J. Vis. Exp. 2019, 2019.

[16] B. C. Hacker, E. J. Lin, D. C. Herman, A. M. Questell, S. E. Martello, R. J. Hedges, A. J. Walker, M. Rafat, Cell. Mol. Bioeng. 2023, 16, 393.

[17] G. M. Sizemore, S. Balakrishnan, K. A. Thies, A. M. Hammer, S. T. Sizemore, A. J. Trimboli, M. C. Cuitiño, S. A. Steck, G. Tozbikian, R. D. Kladney, et al., Nat. Commun. 2018, 9, 2783.

[18] J. P. A. Ioannidis, Sci. Transl. Med. 2012, 4.

[19] R. L. Perlman, Evol. Med. Public Heal. 2016, 2016, 170.

[20] J. Lacombe, F. Zenhausern, Radiother. Oncol. 2022, 176, 187.

[21] N. Gomez-Roman, M. Y. Chong, S. K. Chahal, S. P. Caragher, M. R. Jackson, K. H. Stevenson, S. A. Dongre, A. J. Chalmers, Mol. Cancer Ther. 2020, 19, 575.

[22] C. A. Pasch, P. F. Favreau, A. E. Yueh, C. P. Babiarz, A. A. Gillette, J. T. Sharick, M. R. Karim, K. P. Nickel, A. K. DeZeeuw, C. M. Sprackling, et al., Clin. Cancer Res. 2019, 25, 5376.

[23] D. E. Ingber, Nat. Rev. Genet. 2022, 23, 467.

[24] C. M. Leung, P. de Haan, K. Ronaldson-Bouchard, G. A. Kim, J. Ko, H. S. Rho, Z. Chen, P. Habibovic, N. L. Jeon, S. Takayama, et al., Nat. Rev. Methods Prim. 2022, 2, 33.

[25] S. L. Faley, E. H. Neal, J. X. Wang, A. M. Bosworth, C. M. Weber, K. M. Balotin, E. S. Lippmann, L. M. Bellan, Stem Cell Reports 2019, 12, 474.

[26] V. D. Vest, K. J. Young, I. K. Holtz, M. Mc Veigh, J. D. West, L. M. Bellan, Adv. Healthc. Mater. 2026, 15, e05938.

[27] T. Mathur, K. A. Singh, N. K. R. Pandian, S. H. Tsai, T. W. Hein, A. K. Gaharwar, J. M. Flanagan, A. Jain, Lab Chip 2019, 19, 2500.

[28] N. Gjorevski, B. Avignon, R. Gérard, L. Cabon, A. B. Roth, M. Bscheider, A. Moisan, Lab Chip 2020, 20, 3365.

[29] R. B. Riddle, K. Jennbacken, K. M. Hansson, M. T. Harper, Sci. Rep. 2022, 12, 6855.

[30] M. B. Chen, J. A. Whisler, J. Fröse, C. Yu, Y. Shin, R. D. Kamm, Nat. Protoc. 2017, 12, 865.

[31] R. Riahi, Y. L. Yang, H. Kim, L. Jiang, P. K. Wong, Y. Zohar, Biomicrofluidics 2014, 8.

[32] C. E. Staicu, F. Jipa, I. Porosnicu, A. Bran, E. Stancu, C. Dobrea, B. M. Radu, E. Axente, I. Tiseanu, F. Sima, et al., Appl. Phys. A Mater. Sci. Process. 2022, 128, 770.

[33] L. Y. Yu, C. H. Hsu, C. Y. Li, S. Y. Hong, C. R. Chen, C. S. Chen, Analyst 2023, 148, 3045.

[34] M. Bavoux, Y. Kamio, E. Vigneux-Foley, J. Lafontaine, O. Najyb, E. Refet-Mollof, J. F. Carrier, T. Gervais, P. Wong, Radiother. Oncol. 2021, 157, 175.

[35] S. Jalili-Firoozinezhad, R. Prantil-Baun, A. Jiang, R. Potla, T. Mammoto, J. C. Weaver, T. C. Ferrante, H. J. Kim, J. M. S. Cabral, O. Levy, et al., Cell Death Dis. 2018, 9.

[36] S. Omiya, J. Dalo, Y. Ueda, U. Shankavaram, E. Baldelli, V. Calvert, M. Bylicky, E. F. Petricoin, M. J. Aryankalayil, Radiat. Res. 2025, 203, 293.

[37] S. D. Verma, E. P. de la Chapelle, S. Malkani, C. M. Juran, V. Boyko, S. V. Costes, E. Cekanaviciute, Front. Immunol. 2022, 13, 864923.

[38] Q. Dasgupta, A. Jiang, A. M. Wen, R. J. Mannix, Y. Man, S. Hall, E. Javorsky, D. E. Ingber, Nat. Commun. 2023, 14, 6506.

[39] D. Huh, H. J. Kim, J. P. Fraser, D. E. Shea, M. Khan, A. Bahinski, G. A. Hamilton, D. E. Ingber, Nat. Protoc. 2013, 8, 2135.

[40] B. J. O’Grady, M. D. Geuy, H. Kim, K. M. Balotin, E. R. Allchin, D. C. Florian, N. N. Bute, T. E. Scott, G. B. Lowen, C. M. Fricker, et al., Lab Chip 2021, 21, 4814.

[41] C. Kuperwasser, T. Chavarria, M. Wu, G. Magrane, J. W. Gray, L. Carey, A. Richardson, R. A. Weinberg, Proc. Natl. Acad. Sci. U. S. A. 2004, 101, 4966.

[42] R. G. Bacabac, T. H. Smit, S. C. Cowin, J. J. W. A. Van Loon, F. T. M. Nieuwstadt, R. Heethaar, J. Klein-Nulend, J. Biomech. 2005, 38, 159.

[43] E. Fröhlich, G. Bonstingl, A. Höfler, C. Meindl, G. Leitinger, T. R. Pieber, E. Roblegg, Toxicol. Vitr. 2013, 27, 409.

[44] J. Schindelin, I. Arganda-Carreras, E. Frise, V. Kaynig, M. Longair, T. Pietzsch, S. Preibisch, C. Rueden, S. Saalfeld, B. Schmid, et al., Nat. Methods 2012, 9, 676.

[45] G. Dontu, T. A. Ince, J. Mammary Gland Biol. Neoplasia 2015, 20, 51.

[46] F. Hassiotou, D. Geddes, Clin. Anat. 2013, 26, 29.

[47] K. Ley, C. Laudanna, M. I. Cybulsky, S. Nourshargh, Nat. Rev. Immunol. 2007, 7, 678.

[48] D. Vestweber, Nat. Rev. Immunol. 2015, 15, 692.

[49] X. Wang, Z. Wang, Q. Liao, P. Yuan, J. Mei, Y. Zhang, C. Wu, X. Kang, S. Zheng, C. Yang, et al., Nat. Commun. 2025, 16, 3348.

[50] D. Huh, B. D. Matthews, A. Mammoto, M. Montoya-Zavala, H. Yuan Hsin, D. E. Ingber, Science. 2010, 328, 1662.

[51] Y. Choi, E. Hyun, J. Seo, C. Blundell, H. C. Kim, E. Lee, S. H. Lee, A. Moon, W. K. Moon, D. Huh, Lab Chip 2015, 15, 3350.

[52] D. B. Chou, V. Frismantas, Y. Milton, R. David, P. Pop-Damkov, D. Ferguson, A. MacDonald, Ö. Vargel Bölükbaşı, C. E. Joyce, L. S. Moreira Teixeira, et al., Nat. Biomed. Eng. 2020, 4, 394.

[53] S. Martello, Y. Ueda, M. A. Bylicky, J. Pinney, J. Dalo, K. M. K. Scott, M. J. Aryankalayil, C. N. Coleman, Radiat. Res. 2024, 202, 489.

54. D. H. Han, U. Oh, J. K. Park, ACS Omega 2023, 8, 19128.

[55] P. Parthiban, S. Vijayan, P. S. Doyle, M. Hashimoto, Biomicrofluidics 2021, 15, 024111.

[56] J. Wu, N. Y. Lee, Lab Chip 2014, 14, 1564.

[57] A. M. Malek, S. Izumo, J. Biomech. 1995, 28, 1515.

[58] I. A. Tamargo, K. I. Baek, Y. Kim, C. Park, H. Jo, Nat. Rev. Cardiol. 2023, 20, 738.

[59] G. Follain, D. Herrmann, S. Harlepp, V. Hyenne, N. Osmani, S. C. Warren, P. Timpson, J. G. Goetz, Nat. Rev. Cancer 2020, 20, 107.

[60] G. Follain, N. Osmani, A. S. Azevedo, G. Allio, L. Mercier, M. A. Karreman, G. Solecki, M. J. Garcia Leòn, O. Lefebvre, N. Fekonja, et al., Dev. Cell 2018, 45, 33.

[61] G. Lamberti, B. Prabhakarpandian, C. Garson, A. Smith, K. Pant, B. Wang, M. F. Kiani, Anal. Chem. 2014, 86, 8344.

[62] C. Mondadori, M. Crippa, M. Moretti, C. Candrian, S. Lopa, C. Arrigoni, Front. Bioeng. Biotechnol. 2020, 8, 563471.

[63] S. J. Evani, R. G. Prabhu, V. Gnanaruban, E. A. Finol, A. K. Ramasubramanian, FASEB J. 2013, 27, 3017.

[64] R. Yoshitake, G. Chang, K. Saeki, D. Ha, X. Wu, J. Wang, S. Chen, Front. Cell Dev. Biol. 2022, 10, 850568.

[65] A. D. Reed, S. Pensa, A. Steif, J. Stenning, D. J. Kunz, L. J. Porter, K. Hua, P. He, A. J. Twigger, A. J. Q. Siu, et al., Nat. Genet. 2024, 56, 652.

[66] N. Gjorevski, C. M. Nelson, Nat. Rev. Mol. Cell Biol. 2011, 12, 581.

[67] T. Zhu, S. M. Alves, A. Adamo, X. Wen, K. C. Corn, A. Shostak, S. Johnson, N. D. Shaub, S. E. Martello, B. C. Hacker, et al., Biomaterials 2024, 308, 122531.

[68] M. Vilalta, M. Rafat, A. J. Giaccia, E. E. Graves, Cell Rep. 2014, 8, 402.

[69] G. O. Ahn, J. M. Brown, Cancer Cell 2008, 13, 193.

[70] K. C. Corn, S. E. Martello, V. K. Menon, L. S. Britto, K. M. Simmons, Y. K. Mohamed, Y. I. Ivanova, A. A. Ghelmansaraei, S. A. Weidenbach, T. Zhu, et al., Cell Rep. 2026, 45, 117096.

[71] S. A. C. McDowell, D. F. Quail, Front. Immunol. 2019, 10, 1984.

[72] H. Wijerathne, J. C. Langston, Q. Yang, S. Sun, C. Miyamoto, L. E. Kilpatrick, M. F. Kiani, Radiother. Oncol. 2021, 158, 21.

[73] J. L. Unthank, M. Ortiz, H. Trivedi, L. M. Pelus, C. H. Sampson, R. Sellamuthu, A. Fisher, H. L. Chua, A. Plett, C. M. Orschell, et al., Radiat. Res. 2019, 191, 383.

[74] K. K. C. Tsai, J. Stuart, Y. Y. E. Chuang, J. B. Little, Z. M. Yuan, Radiat. Res. 2009, 172, 306.

[75] J. Severino, R. G. Allen, S. Balin, A. Balin, V. J. Cristofalo, Exp. Cell Res. 2000, 257, 162.

[76] S. A. C. McDowell, R. B. E. Luo, A. Arabzadeh, S. Doré, N. C. Bennett, V. Breton, E. Karimi, M. Rezanejad, R. R. Yang, K. D. Lach, et al., Nat. Cancer 2021, 2, 545.

[77] W. Lee, S. Y. Ko, H. Akasaka, M. Weigert, E. Lengyel, H. Naora, Cancer Cell 2024, 43, 69.

[78] V. Brinkmann, U. Reichard, C. Goosmann, B. Fauler, Y. Uhlemann, D. S. Weiss, Y. Weinrauch, A. Zychlinsky, Science. 2004, 303, 1532.

[79] B. M. Szczerba, F. Castro-Giner, M. Vetter, I. Krol, S. Gkountela, J. Landin, M. C. Scheidmann, C. Donato, R. Scherrer, J. Singer, et al., Nature 2019, 566, 553.

[80] O. Guipaud, C. Jaillet, K. Clément-Colmou, A. François, S. Supiot, F. Milliat, Br. J. Radiol. 2018, 91.

[81] N. Hamada, K. I. Kawano, F. M. Yusoff, K. Furukawa, A. Nakashima, M. Maeda, H. Yasuda, T. Maruhashi, Y. Higashi, Cancers (Basel). 2020, 12, 1.

[82] M. A. Benadjaoud, F. Soysouvanh, G. Tarlet, V. Paget, V. Buard, H. Santos de Andrade, I. Morilla, M. Dos Santos, A. Bertho, B. l’Homme, et al., Int. J. Radiat. Oncol. Biol. Phys. 2022, 112, 975.

[83] X. Luo, Y. Fu, A. J. Loza, B. Murali, K. M. Leahy, M. K. Ruhland, M. Gang, X. Su, A. Zamani, Y. Shi, et al., Cell Rep. 2016, 14, 82.

[84] H. Wijerathne, J. C. Langston, Q. Yang, S. Sun, C. Miyamoto, L. E. Kilpatrick, M. F. Kiani, Radiother. Oncol. 2021, 158, 21.

[85] S. van Kesteren, L. Smeehuijzen, R. Stevenson, J. Kroon, Trends Immunol. 2026, 47, 746.

[86] S. I. Bloom, M. T. Islam, L. A. Lesniewski, A. J. Donato, Nat. Rev. Cardiol. 2023, 20, 38.

[87] O. Guipaud, C. Lago, L. Portier, V. Paget, A. François, S. Supiot, F. Milliat, Br. J. Radiol. 2025, 98, 1176.

[88] H. Yuan, D. J. Goetz, M. W. Gaber, A. C. Issekutz, T. E. Merchant, M. F. Kiani, Radiat. Res. 2005, 163, 544.

[89] D. Hallahan, J. Kuchibhotla, C. Wyble, Cancer Res. 1996, 56, 5150.

[90] J. English, S. Dhanikonda, K. E. Tanaka, W. Koba, G. Eichenbaum, W. L. Yang, C. Guha, JCI Insight 2024, 9.

[91] M. Farjia, C. Pan, Q. Braster, P. Lemnitzer, N. Sachs, L. M. Vöcking, R. Chevre, M. Malamud, C. Schulz, L. Maegdefessel, et al., Arterioscler. Thromb. Vasc. Biol. 2026, 46.

[92] C. A. Reichel, M. Rehberg, M. Lerchenberger, N. Berberich, P. Bihari, A. G. Khandoga, S. Zahler, F. Krombach, Arterioscler. Thromb. Vasc. Biol. 2009, 29, 1787.

[93] H. Wang, L. Chen, C. Wang, Z. Chang, Virulence 2025, 16.

[94] A. K. Gupta, M. B. Joshi, M. Philippova, P. Erne, P. Hasler, S. Hahn, T. J. Resink, FEBS Lett. 2010, 584, 3193.

[95] Z. Guo, C. T. Yang, M. F. Maritz, H. Wu, P. Wilson, M. E. Warkiani, C. C. Chien, I. Kempson, A. R. Aref, B. Thierry, Adv. Mater. Technol. 2019, 4, 1800726.

[96] J. M. Houthuijzen, R. de Bruijn, E. van der Burg, A. P. Drenth, E. Wientjens, T. Filipovic, E. Bullock, C. S. Brambillasca, E. M. Pulver, M. Nieuwland, et al., Nat. Commun. 2023, 14, 183.

