## Supplemental Figures S1-S4 and Table S1 for "A mammary-specific microfluidic device for studying post-radiotherapy vascular-immune cell interactions"

**Supplementary Material**

**Supplemental Table S1. Mouse Cytokine Array Analytes.**

| **Name** | **Abbreviation** |
| --- | --- |
| Granulocyte-macrophage colony-stimulating factor | GM-CSF |
| Interferon gamma | IFNγ |
| Interleukin-1 beta | IL-1β |
| Interleukin-2 | IL-2 |
| Interleukin-4 | IL-4 |
| Interleukin-5 | IL-5 |
| Interleukin-6 | IL-6 |
| Interleukin-10 | IL-10 |
| Interleukin-12 p70 | IL-12p70 |
| Interleukin-13 | IL-13 |
| Interleukin-17A | IL-17A |
| Interleukin-17F | IL-17F |
| Monocyte chemoattractant protein-1/C-C motif chemokine ligand 2 | MCP-1/CCL2 |
| Tumor necrosis factor alpha | TNF⍺ |

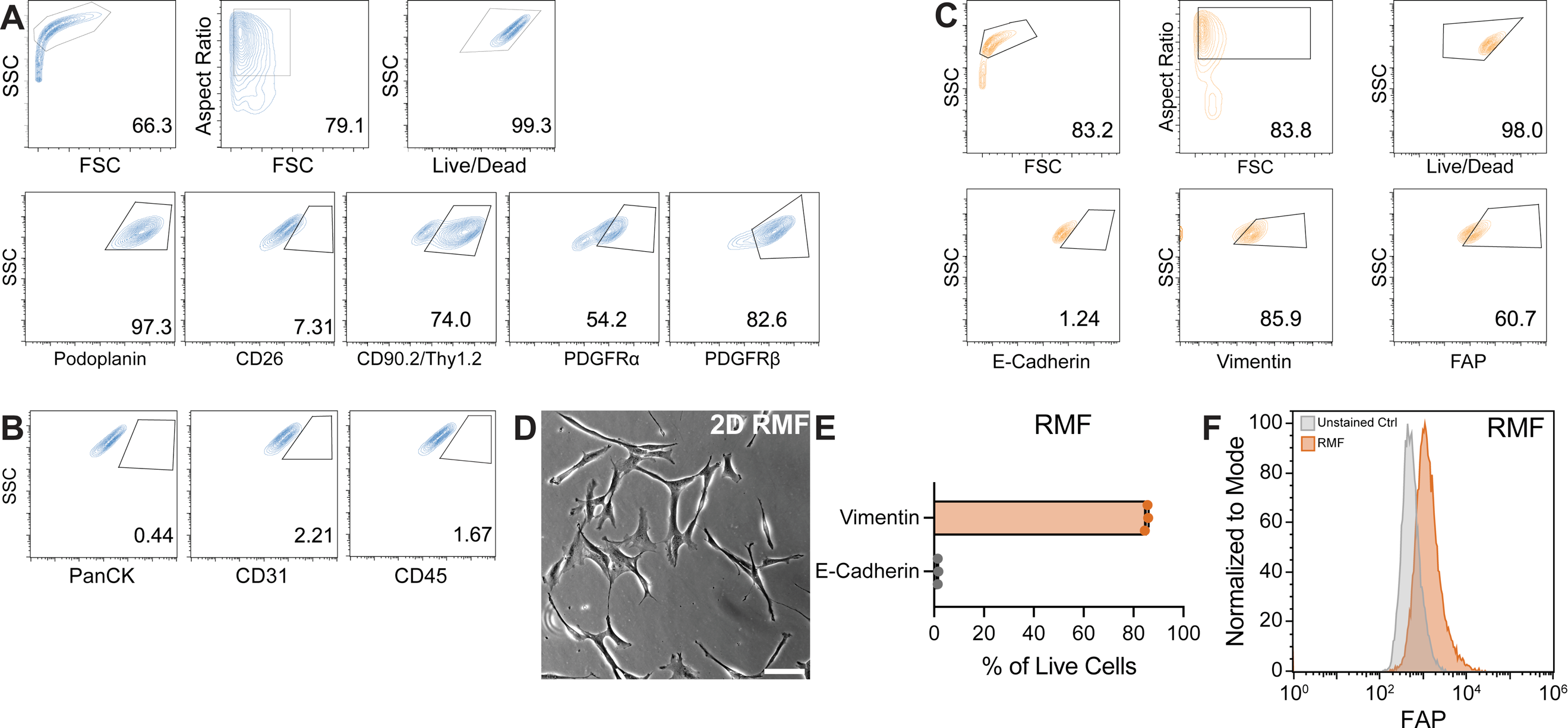
**Supplemental Figure S1. SVF and RMF flow cytometry analysis. A.** Gating strategy for SVF analysis. **B.** Negligible expression of PanCK, CD45, and CD31 in SVF cells before adding to MVoCs. **C.** Gating strategy for RMF analysis. **D.** Representative phase contrast image of RMF cells in 2D culture. Scale bar is 100 µm. **E.** Flow cytometry characterization of RMFs prior to seeding in MVoCs. n=3 independent experiments. **F.** Representative flow cytometry histogram analysis of fibroblast activation protein (FAP) expression in RMFs prior to seeding in MVoCs.

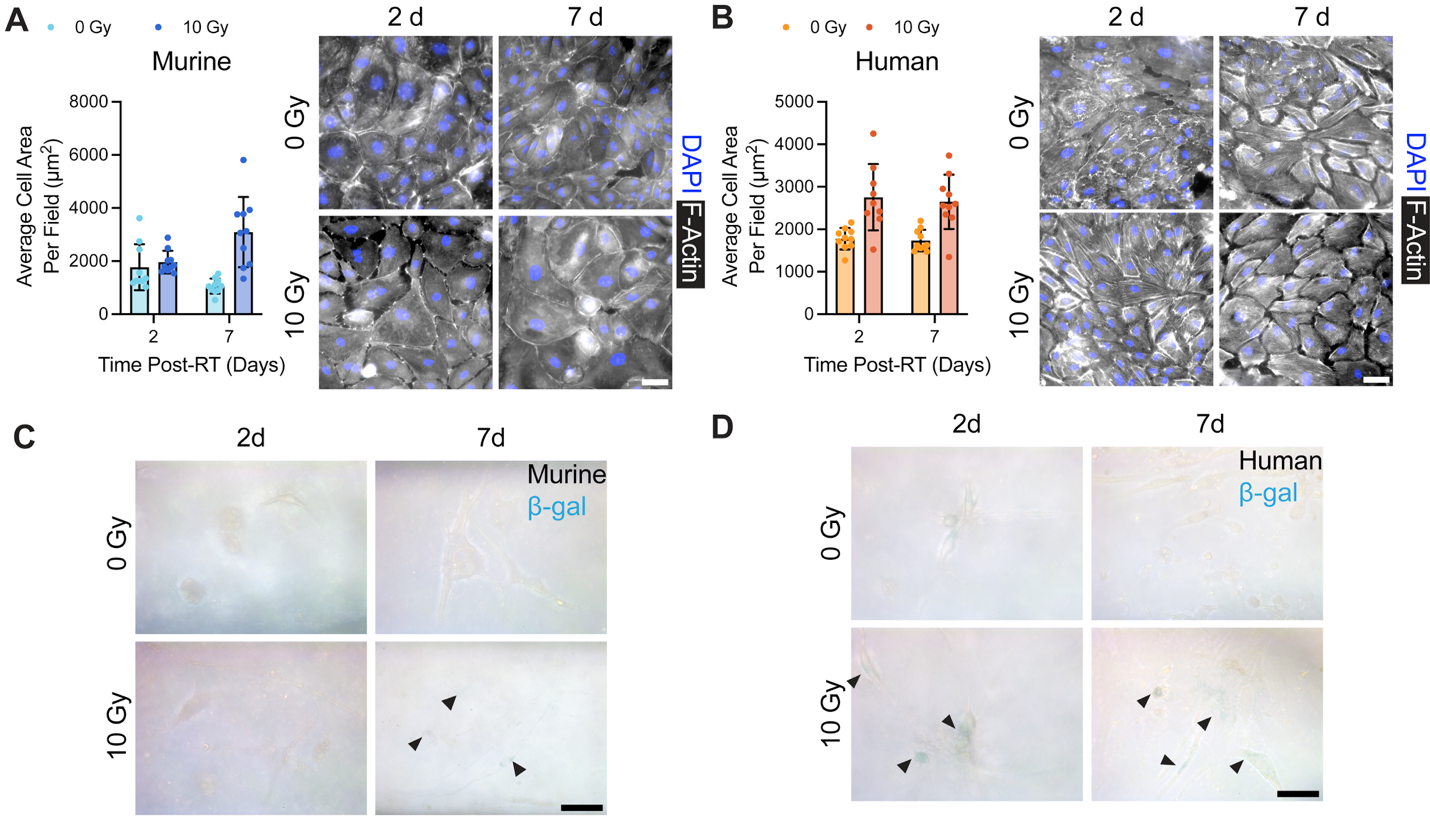
**Supplemental Figure S2. Endothelial cell size and mammary fibroblast-like beta-galactosidase (β-gal) activity increase post-RT.** Quantification and representative images of murine **(A)** and human **(B)** cell area post-RT determined by F-Actin staining of EC monolayers. Each point is one representative image with 10 images per condition. β-gal activity staining in mouse SVF cells **(C)** and human RMFs **(D)**. Arrows indicate positive staining. Scale bars are 50 µm.

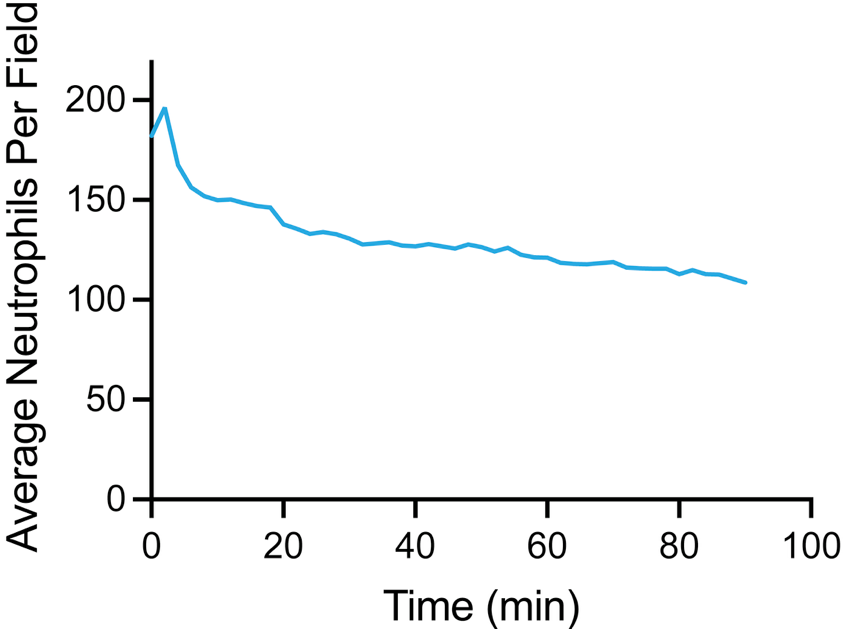

**Supplemental Figure S3. Neutrophil adhesion over time.** Murine MVoCs were irradiated to 10 Gy and cultured for 48 h. Deep Red CellTracker-stained neutrophils were then perfused through the devices and images were captured every 2 min for 90 min total. Five points along the device channel were imaged at each timepoint and the number of neutrophils at each point was averaged to obtain the average number of neutrophils per field over time.

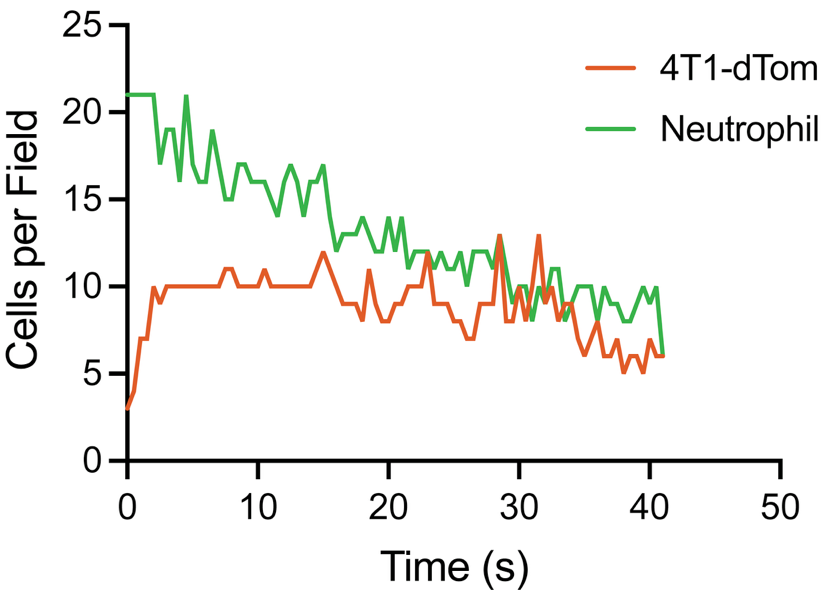

**Supplemental Figure S4. Tumor cell and neutrophil adhesion over time.** Murine MVoCs were irradiated to 10 Gy and cultured for 48 h before perfusing Deep Red CellTracker-stained neutrophils for 30 min. 4T1-dTomato (4T1-dTom) cells were then perfused for perfused through the devices and images were captured every 30 s for 45 min total. Five points along the device channel were imaged at each timepoint and the number of neutrophils and 4T1-dTom at each point was averaged to obtain the number of cells per field.

**Supplemental Video S1. Tumor cell and neutrophil adhesion over time.** Live cell imaging (20X) of 4T1-dTom (red) adhesion post-neutrophil adhesion (green) over 45 min. Each frame is one image taken every 30 s.
